# Mitochondrial membrane junction-mediated ATP channeling drives activity-dependent glucose metabolism

**DOI:** 10.64898/2025.12.18.695286

**Authors:** Dengbao Yang, Gemma Molinaro, Nadine Nijem, Chuanhai Zhang, Dylen Penny, Meijuan Bai, Mei-Jung Lin, Xiaodong Wen, Salvador Pena, Janaka Wansapura, Maria Gaitanou, Rebecca Matsas, Boyuan Wang, Hamid Baniasadi, Yan Han, Jay Gibson, Kimberly Huber, Xing Zeng

## Abstract

Neurons and brown adipocytes rely on rapid ATP production from accelerated glucose metabolism to sustain bursts of activity upon stimulation, a process known as activity-dependent glucose metabolism. The first committed step in this pathway, the hexokinase I (HK1)-catalyzed phosphorylation of glucose, consumes ATP, raising the question of how this reaction can be accelerated when cytosolic ATP becomes limiting during stimulation. We identify Cell Cycle Exit and Neuronal Differentiation protein 1 (CEND1), expressed in both cell types, as a critical regulator of this process. Loss of CEND1 impairs activity-dependent glucose utilization, ATP generation, and stimulation-evoked activity both *in vitro* and *in vivo*. Mechanistically, CEND1 assembles a complex with HK1, voltage-dependent anion channel 1 (VDAC1), and adenine nucleotide translocase 1 (ANT1) at hemifusion-like membrane junction between the outer/inner mitochondrial membrane, channeling mitochondrially derived ATP directly to HK1. These findings uncover a previously unrecognized mechanism that sustains activity-dependent glucose metabolism, with broad implications for energy homeostasis in specialized cell types.

## Introduction

Cells tailor their energy metabolism to differentiation state and level of activity. Neurons and brown adipocytes exemplify this principle, as they constantly switch from resting to active stimulated states with sharply elevated energy demands. In neurons, action potential firing and synaptic transmission require large amounts of ATP to sustain vesicle cycling, neurotransmitter loading, and resetting cross-membrane ion gradients^1,2^. In brown adipocytes, adrenergic stimulation drives ATP consumption through cAMP synthesis and fatty acyl-CoA production^3^. In both cell types, glucose utilization rises rapidly during activation^4–7^, and inhibition of glucose metabolism compromises their stimulus-evoked functions^2,8–14^. Impaired cerebral glucose metabolism is a hallmark of aging and Alzheimer’s disease (AD), preceding symptoms and correlating with progression^15–19^, whereas glucose administration can transiently improve cognition in older adults and AD patients^20–23^, underscoring the link between metabolic dysfunction and cognitive decline. These observations highlight the importance of glucose metabolism in meeting activity-driven energy needs in diverse cellular contexts and raise a central question: how do neurons and brown adipocytes accelerate glucose metabolism and ATP production upon activation?

In neurons, activity triggers the translocation of the glucose transporter GLUT4 to axonal membranes, thereby enhancing glucose influx^24^. However, a key metabolic bottleneck arises downstream. Once inside the cell, glucose must be phosphorylated to glucose-6-phosphate (G6P) in an ATP-dependent reaction catalyzed by hexokinases, the first rate-limiting step of glucose metabolism that traps glucose intracellularly. During periods of high activity, when ATP is rapidly consumed by diverse processes, it is unclear how the hexokinase reaction could be accelerated to promote glucose metabolism. One plausible solution is that hexokinases access a privileged source of ATP, protected from direct competition with other ATP-demanding reactions. Supporting this idea, the predominant neuronal isoform, hexokinase I (HK1), is anchored to the outer mitochondrial membrane (OMM)^25^, mediated by its N-terminal amphipathic motif, interaction with voltage-dependent anion channels (VDACs), and O-GlcNAcylation^26–29^. VDACs mediate adenine nucleotide exchange across the OMM^30,31^, and *in vitro* assays demonstrate that VDAC-bound HK1 preferentially utilizes mitochondrially generated ATP^32^. This supports a model in which mitochondrial ATP export is directly coupled to HK1 activity, ensuring continuous glucose phosphorylation even when cytosolic ATP is limiting. Nevertheless, a critical gap remains for this model: ATP synthesized in the mitochondrial matrix must first be exported into the intermembrane space (IMS) by the adenine nucleotide translocator (ANT) in the inner mitochondrial membrane (IMM), from which it is thought to diffuse freely. How ATP is preferentially funneled to VDAC-bound HK1, rather than dispersing through VDACs unoccupied by HK1, remains unknown. Such directional channeling could be essential for the tight coupling of mitochondrial ATP production to HK1 activity. Whether this mechanism exists, and how it might be achieved at a molecular level, has not been explored. However, given the shared capacity of neurons and brown adipocytes to rapidly upregulate glucose metabolism during activation, it is intriguing to postulate that both cell types may have convergently evolved a similar strategy to ensure efficient glucose phosphorylation despite their developmental and functional distinctions.

Cell cycle exit and neuronal differentiation protein 1 (CEND1) is a neural-specific, single-pass transmembrane protein, first identified as the target of the monoclonal antibody BM88 in brain membrane fractions across multiple vertebrate species^33^. Its expression begins in the late stage of neural progenitors, peaks during terminal differentiation, and declines with age^34,35^. Early studies of CEND1-deficient mice suggested mild neurodevelopmental abnormalities^36^, but more recent work has revealed a broader role in cognition. CEND1 deficiency impairs learning and memory in mice^37^. CEND1 levels are reduced in Alzheimer’s disease (AD) models, and CEND1 overexpression enhances memory performance^38^. The *Cend1* transcript is enriched in neurons activated during memory-related tasks^39^, and human genetic studies link CEND1 expression to cognitive resilience in both aging and depression^40,41^. Strikingly, the only common genetic variant significantly associated with circulating CEND1 levels in the UK Biobank (>54,000 participants) is ApoE-ε4, the strongest genetic risk factor for late-onset AD^42^. Collectively, these findings establish CEND1 as a critical contributor to cognitive function, although its molecular mechanism of action remains unknown.

Here, we show that CEND1 is highly expressed in both neurons and cold-stimulated brown adipocytes, where it is essential for activity-dependent increases in glucose metabolism. CEND1 loss impairs glucose utilization, ATP production, and stimulus-evoked functional responses in both cell types. Mechanistically, CEND1 assembles into a complex with HK1, VDAC1, and ANT1 at membrane junctions connecting OMM and IMM, enabling direct channeling of mitochondrial ATP to HK1 and sustaining G6P formation during high energy demand. These findings reveal a previously unrecognized mechanism for targeted ATP delivery across mitochondrial membranes, representing a new principle of metabolic coupling with broad physiological significance for both excitable and thermogenic cells.

## Results

### CEND1 contributes to brown adipocyte thermogenesis

While investigating the role of the histone demethylase LSD1 in brown adipocytes, we identified *Cend1* as a direct transcriptional target of LSD1-mediated repression (Figure S1A-B)^43^. Tissue profiling of mice housed at room temperature (23 °C) showed that *Cend1* expression is most abundant in the brain and brown adipose tissue (BAT) (Figure 1A). Analysis of published translating ribosome affinity purification sequencing (TRAP-seq) datasets driven by adipocyte-specific Cre recombinase confirmed higher expression of *Cend1* in brown relative to beige and white adipocytes (Figure S1C-D)^44,45^. Notably, despite strong LSD1 repression, both *Cend1* mRNA and protein levels were markedly elevated in the BAT from cold-acclimated mice (4 °C) compared with those housed at thermoneutrality (30 °C) (Figure 1B-C), suggesting a role for CEND1 in cold-induced BAT thermogenesis.

**Figure 1.**
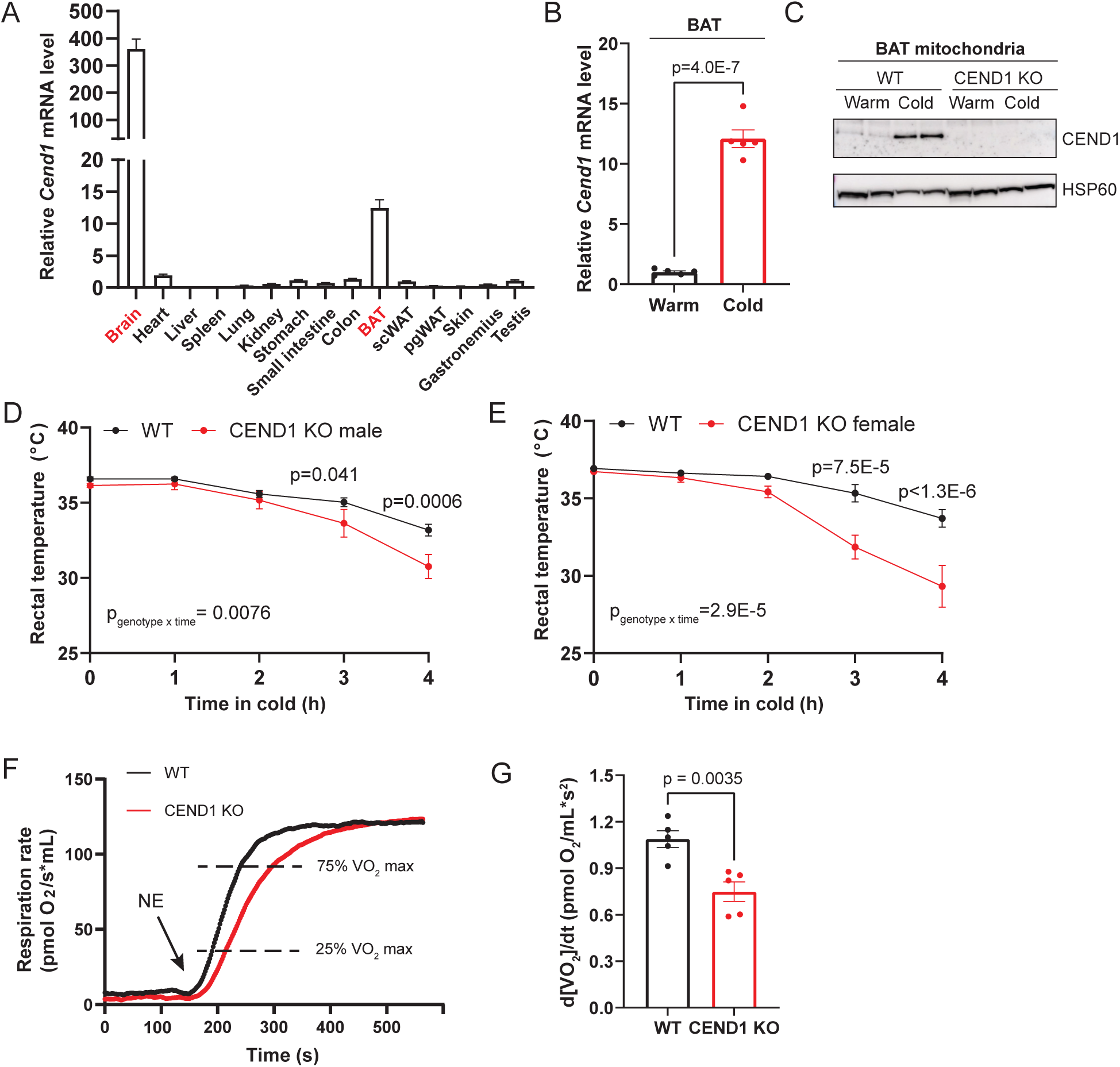
CEND1 contributes to brown adipocyte thermogenesis. A. qPCR analysis of *Cend1* mRNA level across mouse tissue (*n* = 3). B. qPCR analysis of *Cend1* mRNA levels in the BAT from warm or cold-acclimated mice (*n* = 5 per group). C. Western blot analysis of CEND1 protein levels in the BAT mitochondria isolated from warm or cold-acclimated WT or CEND1 KO mice. D-E. Cold tolerance test of male (D) and female (E) WT and CEND1 KO mice (*n* = 10 per group for D; *n* = 8 per group for E). F. Representative oxygen consumption traces of 100,000 freshly isolated WT and CEND1 KO brown adipocytes. G. Time derivative of oxygen consumption rates (VO_2_) of WT and CEND1 KO brown adipocytes within the 25%-75% VO_2_max interval (*n* = 5 per group). Data represent mean ± s.e.m. Statistical significance was calculated by two-tailed unpaired t-test (B, F) or two-way ANOVA with repeated measurements (D, E).

To test this possibility, we performed cold tolerance assays in wild-type (WT) and *Cend1* knockout (KO) mice. Both male and female KO mice displayed heightened cold sensitivity relative to WT controls (Figure 1D-E), consistent with impaired BAT thermogenic function. Because CEND1 is also expressed in sympathetic neurons (Figure S1E and Furlan et al^46^), which is essential to drive BAT thermogenesis, we asked whether defective sympathetic activity might account for this phenotype. We found that norepinephrine (NE) turnover rates, an indicator of sympathetic nerve activity, were comparable in WT and KO BAT during acute cold exposure (Figure S1F). Thus, impaired sympathetic drive is unlikely to be a major contributor to the cold intolerance.

To further exclude non-cell-autonomous effects, we assessed the respiratory activity of primary brown adipocytes freshly isolated from WT and KO mice. Upon NE stimulation *in vitro*, CEND1-deficient brown adipocytes exhibited consistently slower respiratory responses, as measured by the time derivative of the oxygen consumption rates (VO_2_) within the interval 25%-75% VO_2_max, during which VO_2_ increases approximately linearly with time (Figure 1F-G). These findings confirmed a cell-autonomous role for CEND1 in thermogenic response.

### CEND1 promotes brown adipocyte activity-dependent glycolysis and ATP production

To investigate the mechanism underlying the blunted thermogenic response of *Cend1* KO brown adipocytes, we examined the adrenergic receptor-cAMP-PKA signaling axis, which drives thermogenic activation downstream of NE stimulation. We first asked whether cAMP production was altered in the absence of CEND1. Upon NE stimulation, *Cend1* KO brown adipocytes exhibited significantly reduced cAMP levels compared to WT controls (Figure 2A), a difference more clearly demonstrated in the presence of the phosphodiesterase inhibitor IBMX (Figure 2A), reflecting the short half-life of cAMP. Given the high ATP demand required to support rapid turnover of cAMP in brown adipocytes, we hypothesized that the reduced cAMP levels in *Cend1* KO cells reflect impaired ATP production.

**Figure 2.**
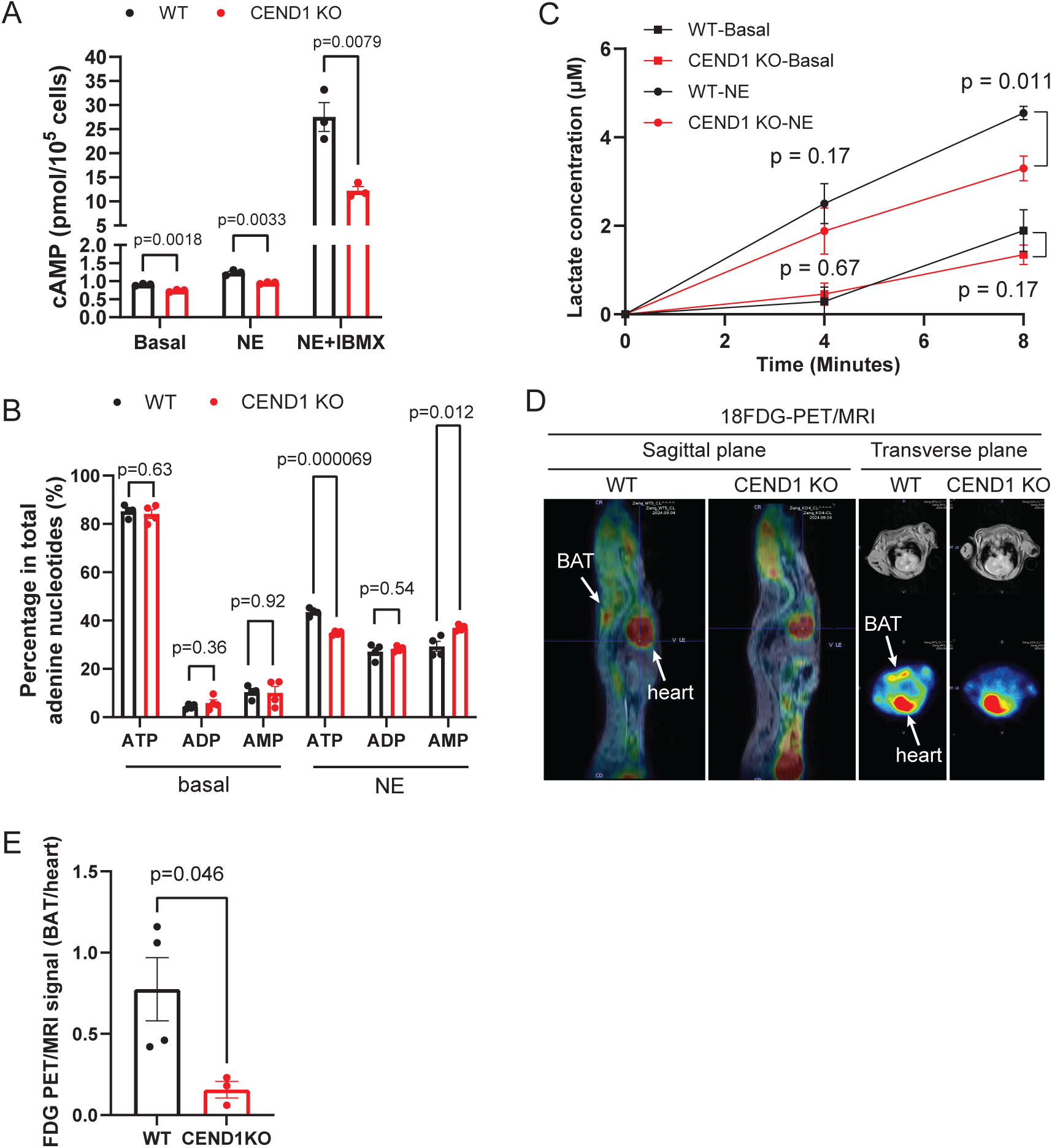
CEND1 promotes brown adipocyte activity-dependent glycolysis and ATP production. A. Quantitation of cAMP levels in WT or CEND1 KO brown adipocytes with or without NE stimulation (*n* = 3 per group). B. Quantitation of adenine nucleotide levels in WT or CEND1 KO brown adipocytes with or without NE stimulation (*n* = 4 per group). C. Quantitation of lactate released from WT or CEND1 KO brown adipocytes with or without NE stimulation (*n* = 4 per group). D-E. Representative ^18^FDG-PET/MRI images (D) and quantitation (E) of WT or CEND1 KO mice treated with the β3-specific adrenergic receptor agonist CL316,243 (*n* = 4 for WT; *n* = 3 for *Cend1* KO). Data represent mean ± s.e.m. Statistical significance was calculated by two-tailed unpaired t-test (A, B, and E) or two-way ANOVA with repeated measurements (C).

To test this, we quantified ATP, ADP, and AMP levels in WT and KO brown adipocytes under basal and NE-stimulated conditions using a luminescence approach combined with enzymes interconverting adenine nucleotides (Figure S2A-E). At baseline, ATP was the predominant adenine nucleotide in both genotypes (WT 85%, CEND1 KO 84% as a fraction in total adenine nucleotides, Figure 2B). NE stimulation triggered substantial reductions in ATP levels in both groups, with the KO cells exhibiting more profoundly decreased ATP levels (WT 44% remaining, KO 35% remaining), accompanied by significantly higher AMP levels than the WT cells (WT 29%, KO 37%, Figure 2B). These findings indicate that CEND1 is required to sustain ATP production in response to NE, thereby supporting cAMP synthesis and activation of the downstream thermogenic response.

Because brown adipocytes have limited oxidative phosphorylation capacity, owing to UCP1-mediated uncoupling and inefficient ATP synthase assembly^47,48^, they rely heavily on glycolysis for ATP production during activation^9,10^. We therefore tested whether *Cend1* KO brown adipocytes exhibit defective glycolytic responses. Lactate release, a proxy for glycolytic flux, was similar between genotypes under basal conditions (Figure 2C). Upon NE stimulation, lactate release increased in both groups, but KO cells produced significantly less lactate than WT (Figure 2C), consistent with impaired glycolytic upregulation.

To validate these findings *in vivo*, we performed positron emission tomography-magnetic resonance imaging (PET-MRI) analysis in WT and *Cend1* KO mice treated with the β3-adrenergic receptor agonist CL316,243. This direct adipocyte-specific pharmacologic stimulation bypasses the sympathetic nervous system, eliminating potential confounds from CEND1-related neuronal deficits. Under these conditions, *Cend1* KO mice exhibited significantly reduced ^18^F- fluorodeoxyglucose (FDG) accumulation in the BAT compared to WT (Figure 2D-E), further confirming that CEND1 is essential for activity-dependent glucose metabolism in brown adipocytes.

### CEND1 promotes neuronal activity-dependent glucose metabolism

Similar to brown adipocytes, neurons also display activity-dependent increases in glucose metabolism, which is important to support energy-demanding processes, such as maintaining membrane excitability and synaptic vesicle cycling. We therefore asked whether CEND1 plays a similar role in promoting activity-dependent glucose metabolism in neurons. For this purpose, we set up a system for simultaneously measuring real time glucose level changes and neuronal electrical activity in response to stimulation in WT and *Cend1* KO hippocampal slices (Figure 3A). In some slices, we recorded population spikes after field excitatory postsynaptic potential (fEPSP) (Figure 3B), reflecting synchronized action potential (AP) firing from groups of neurons^49^. Compared with slices in which only fEPSP was recorded, the glucose sensor recorded significantly more profound local glucose depletion in WT slices with population spikes (Figure 3B-C), reflecting activity-dependent increases glucose utilization. In contrast, the glucose responses from KO slices with population spikes were substantially attenuated compared with the WT slices (Figure 3B-C). These data thus show that CEND1 deficiency impairs neuronal activity-dependent glucose utilization.

**Figure 3.**
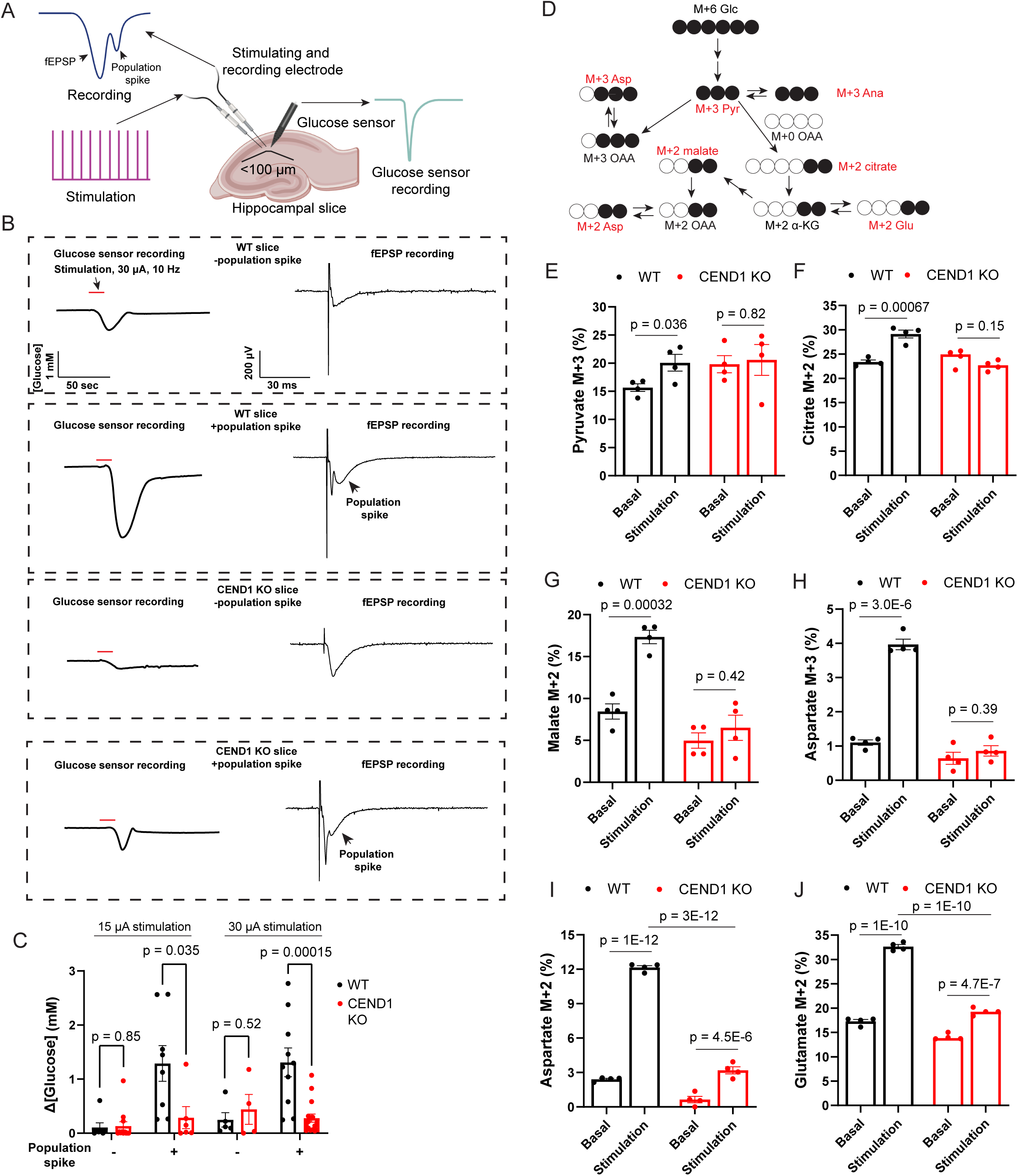
CEND1 is required for neuronal activity-dependent glucose metabolism. A. Schematics of the electrical stimulation/glucose biosensor recording setup. B-C. Representative traces from the recording electrode and glucose biosensor (B) and quantitation of the responses of WT or CEND1 KO slices to electrical stimulation (C) (*n* = 4-12 for no population spike group; *n* = 6-16 for population spike + group). The first responses of the 150 pulse train are shown in the traces. The scale bars and stimulation mark apply to all traces. D. Schematics of ^13^C-glucose labeling of glycolysis and TCA cycle intermediates and transamination products. E-J. ^13^C labeling efficiency of various metabolites in DIV11 primary WT and CEND1 KO cortical neurons with or without electrical stimulation (*n* = 4 per group). Data represent mean ± s.e.m. Statistical significance was calculated by two-tailed unpaired t-test (C, E, F, G, and H) or two-way ANOVA (I and J).

To demonstrate this point using an alternative strategy, we performed isotope tracing in primary cortical neurons cultured *in vitro*. CEND1 expression was found to gradually increase as those neurons mature (Figure S3A-B). On Day *in vitro* (DIV) 11, we switched neurons cultured in the regular medium containing ^12^C-glucose to the same medium containing U-^13^C-glucose and harvested them after 10 minutes with or without electrical stimulation during the first minute (10 Hz, 1 minute duration). For WT neurons, the fraction of ^13^C-labeled pyruvate M+3, TCA cycle intermediates citrate M+2 and malate M+2, aspartate M+3 (transamination/carboxylation product of pyruvate M+3, Figure 3D), all increased significantly from the no stimulation to the stimulation condition (Figure 3E-H), reflecting activity-dependent increases in glucose-derived carbon fluxes. In striking contrast, no significant activity-dependent increase of pyruvate M+3, citrate M+2, malate M+2, and aspartate M+3 labeling was observed in KO neurons (Figure 3E-H). Pronounced activity-dependent labeling of glutamate (glutamate M+2, transamination product of α-ketoglutarate M+2) and aspartate (aspartate M+2, transamination product of oxaloacetate M+2) were also observed in WT neurons, suggesting high fluxes through the malate aspartate shuttle (MAS) responsible for shuttling glycolysis-derived NADH equivalents to mitochondria and sustaining glycolytic flux (Figure 3I-J). In contrast, the magnitude of glutamate and aspartate labeling was substantially attenuated in the KO, consistent with reduced MAS and glycolytic fluxes (Figure 3I-J). Lactate labeling did not reveal a significant activity-dependent increase in either genotype, suggesting that activity-induced glucose metabolism predominantly proceeds through oxidative phosphorylation (Figure S3C). Taken together, these data support a critical role for CEND1 in neuronal activity-dependent glucose metabolism, consistent with its function in brown adipocytes.

Notably, under resting conditions, KO neurons showed a trend toward increased ^13^C labeling of both pyruvate and lactate, significant by unpaired t-test but not by the more stringent two-way ANOVA (Figure S3C-D), without corresponding increases in TCA cycle intermediates or transamination products (Figure 3E-J). These findings suggest that CEND1 deficiency enhances resting-state glycolytic flux without proportionally increasing TCA cycle activity, a point which we will develop with additional observations.

### CEND1 deficiency impairs neuronal activity-dependent ATP production, activity and increases seizure threshold

Neuronal activity-dependent glucose utilization is essential for supporting the rapid ATP demands triggered by stimulation. To evaluate whether CEND1 contributes to this process, we quantified ATP levels in primary WT and *Cend1* KO cortical neurons cultured *in vitro* following brief electrical stimulation (10 Hz, 1 min). The KO neurons exhibited a much more profound reduction in ATP levels than the WT neurons after stimulation (Figure 4A-B, 60% reduced for KO and 28% for WT). These findings show that CEND1 deficiency impairs activity-dependent ATP production in neurons. Interestingly, we observed that total adenine nucleotide levels also dropped markedly after stimulation in both genotypes, and the magnitude of reduction was significantly greater in KO neurons than in WT neurons (Figure 4C-D, 43% for KO vs. 17% for WT). This pronounced loss of total adenine nucleotides in neurons suggested conversion into alternative metabolites. Prior studies have shown that AMP deaminase is a major route for AMP disposal during neuronal activity, producing IMP (Figure 4E)^50^, which may prevent excessive AMP accumulation and further depletion of ATP through the adenylate kinase reaction. Consistent with this, we observed substantially higher activity-dependent IMP accumulation in KO neurons compared with WT neurons (Figure 4F), accounting for their larger decline in total adenine nucleotide levels. A similar but more modest reductions in total adenine nucleotides level after NE stimulation was also observed in brown adipocytes (WT 9%, KO 16%, Figure S4), supporting more activity-driven AMP-IMP conversion in the KO brown adipocytes, which was also noted in a prior study^51^. Together, these results support a model in which CEND1 is required for efficient ATP production during neuronal activation, paralleling its role in brown adipocytes.

**Figure 4.**
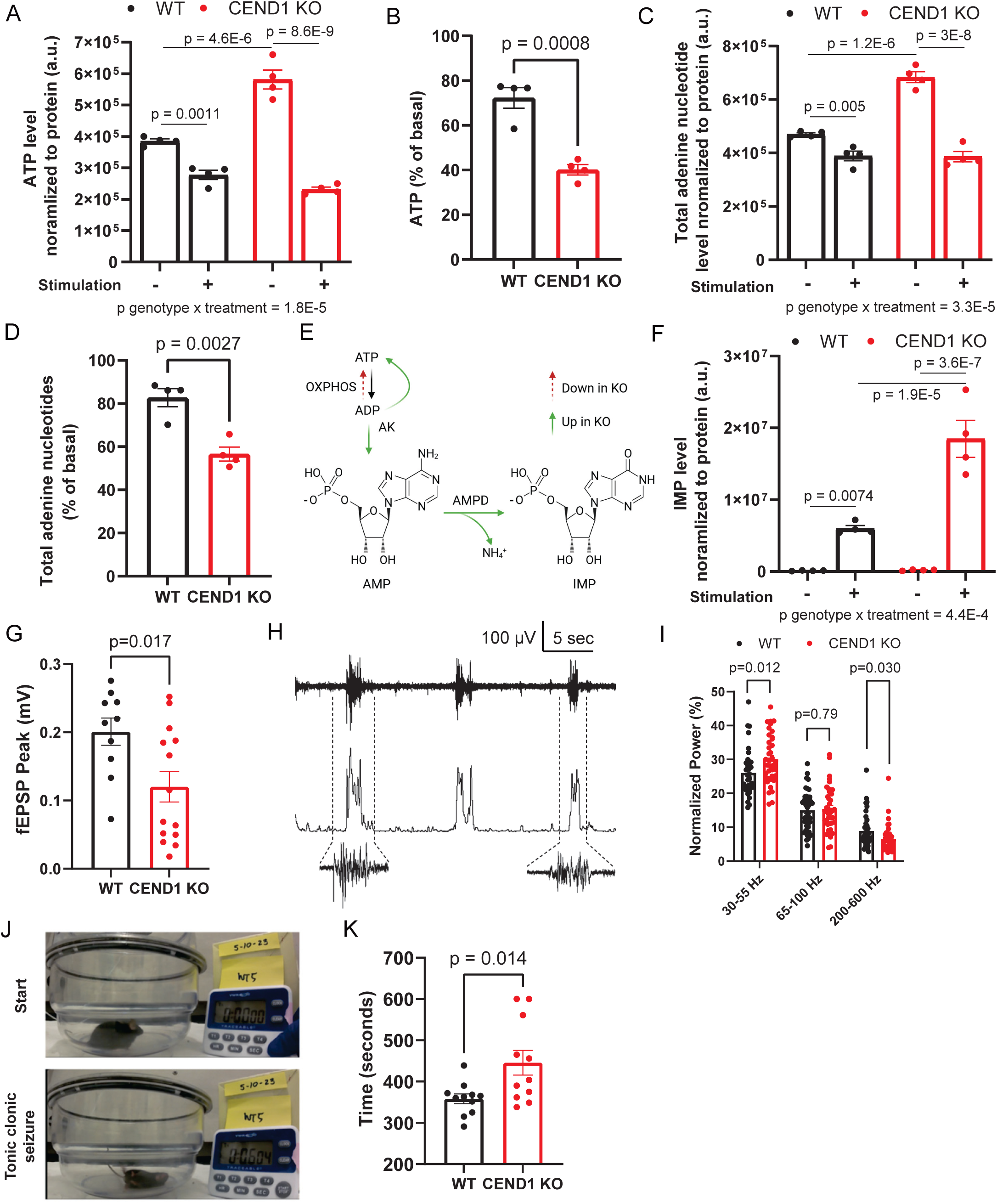
CEND1 deficiency impairs neuronal activity-dependent ATP production, activity and increases seizure threshold. A. ATP levels normalized to protein in DIV11 primary WT and CEND1 KO cortical neurons with or without electrical stimulation (*n* = 4 per group). B. Stimulation-induced relative reduction in ATP in DIV11 primary WT and CEND1 KO cortical neurons with or without electrical stimulation (*n* = 4 per group). C. Total adenine nucleotides levels in DIV11 primary WT and CEND1 KO cortical neurons with or without electrical stimulation (*n* = 4 per group). D. Stimulation-induced relative reduction in total adenine nucleotides in DIV11 primary WT and CEND1 KO cortical neurons (*n* = 4 per group). E. Illustration of the interconversion between nucleotides. AK, adenylate kinase. AMPD, AMP deaminase. Note that in KO neurons, deficiency in ATP regeneration via oxidative phosphorylation results in more AMP production by adenylate kinase, which is subsequently converted to IMP. F. IMP levels normalized to protein in DIV11 primary WT and CEND1 KO cortical neurons with or without electrical stimulation (*n* = 4 per group). G. fEPSP peak responses of WT and CEND1 KO slices stimulated at 8 μA, 10 Hz (*n* = 10 for WT; *n* = 14 for *Cend1* KO). H. Representative trace of UP states recorded in a slice. I. Normalized power measured during UP states for 3 frequency bands, 200-600 Hz is decreased while 30-55 Hz is increased (*n* = 37-40 per group). J. Mouse postures at the start or the onset of tonic-clonic seizure in the fluorothyl exposure experiment. K. Latency from initiation of flurothyl exposure to onset of tonic-clonic seizures (*n* = 11 per group). Data represent mean ± s.e.m. Statistical significance was calculated by two-way ANOVA (A, C and F), unpaired t-test (B, D, G, and K), or Mann-Whitney t-tests with Bonferroni correction (I).

Interestingly, under unstimulated conditions, KO neurons exhibited significantly higher levels of ATP and total adenine nucleotides than the WT (Figure 4A and C). This observation is consistent with the enhanced ^13^C-pyruvate and lactate labeling in KO neurons at rest (Figure S3C-D), suggesting enhanced resting state glucose metabolism and likely nucleotide synthesis driven by the pentose phosphate pathway in the absence of CEND1.

Reduced activity-dependent ATP production is predicted to lead to impaired neuronal activity. This is supported by multiple lines of observations. First, *Cend1* KO hippocampal slices displayed significantly reduced fEPSP peak values than the WT at a stimulation intensity not sufficient to trigger population spikes (8 μA, 10 Hz, 150 stimulations, Figure 4G), indicating impaired synaptic transmission. Second, when we examined spontaneously occurring activity states in cortical slices (often referred to as UP states^52–54^), we found the normalized power in *Cend1* KO slices was altered relative to WT – decreased in the 200-600 Hz band and increased in the 30-55 Hz band (Figure 4H-I). Given that signals originating from synaptic transmission are observed at frequencies <100 Hz and signals from APs are mainly at >200 Hz^55^, these results suggest less AP firing relative to synaptic signaling in the *Cend1* KO brain slices and support that CEND1 is important for neural activity. Lastly, given the mounting evidence that glucose restriction has anti-epileptic effects, we analyzed the contribution of CEND1 to seizure sensitivity. We subjected WT and *Cend1* KO mice to flurothyl treatment, a volatile chemoconvulsant acting as a GABAα antagonist previously used to induce seizures in both humans and animals. Upon exposure to flurothyl, *Cend1* KO mice exhibited significantly delayed onset of tonic-clonic seizure than WT mice (Figure 4J-K), suggesting that ablation of CEND1 raises the seizure threshold, consistent with impaired glucose usage in KO neurons.

Taken together, our results show that CEND1 is crucial for neuronal activity by promoting neuronal activity-dependent glucose metabolism and ATP production, a role shared with brown adipocytes, despite two cell types being developmentally and functionally distinct.

### CEND1 forms a complex with HK1/VDAC1/ANT1 at OMM/IMM junction

Our analyses revealed that CEND1 promotes activity-dependent glucose metabolism in both neurons and brown adipocytes, suggesting a shared mechanism across these distinct cell types. To explore this mechanism, we first examined CEND1’s subcellular localization. Previous studies showed that CEND1 is a single-pass transmembrane protein localized to the outer mitochondrial membrane (OMM) in neurons^38^, which we confirmed in both central and peripheral neurons (Figure S1E and S3B). Using immunofluorescence and proteinase K digestion, we further demonstrated that endogenous CEND1 also localizes to the OMM in brown adipocytes (Figure 5A-B). Similarly, exogenously expressed CEND1 in HeLa cells exhibited OMM localization (Figure S5A). Mutagenesis analysis revealed that substituting the polybasic C-terminal RKK motif with GGG or replacing evolutionarily conserved glycine residues in the transmembrane domain with bulky tryptophan residues (G129G130 to WW) resulted in CEND1 mislocalization (Figure S5B-C). These findings indicate that CEND1 belongs to the family of tail-anchored OMM-targeting proteins.

**Figure 5.**
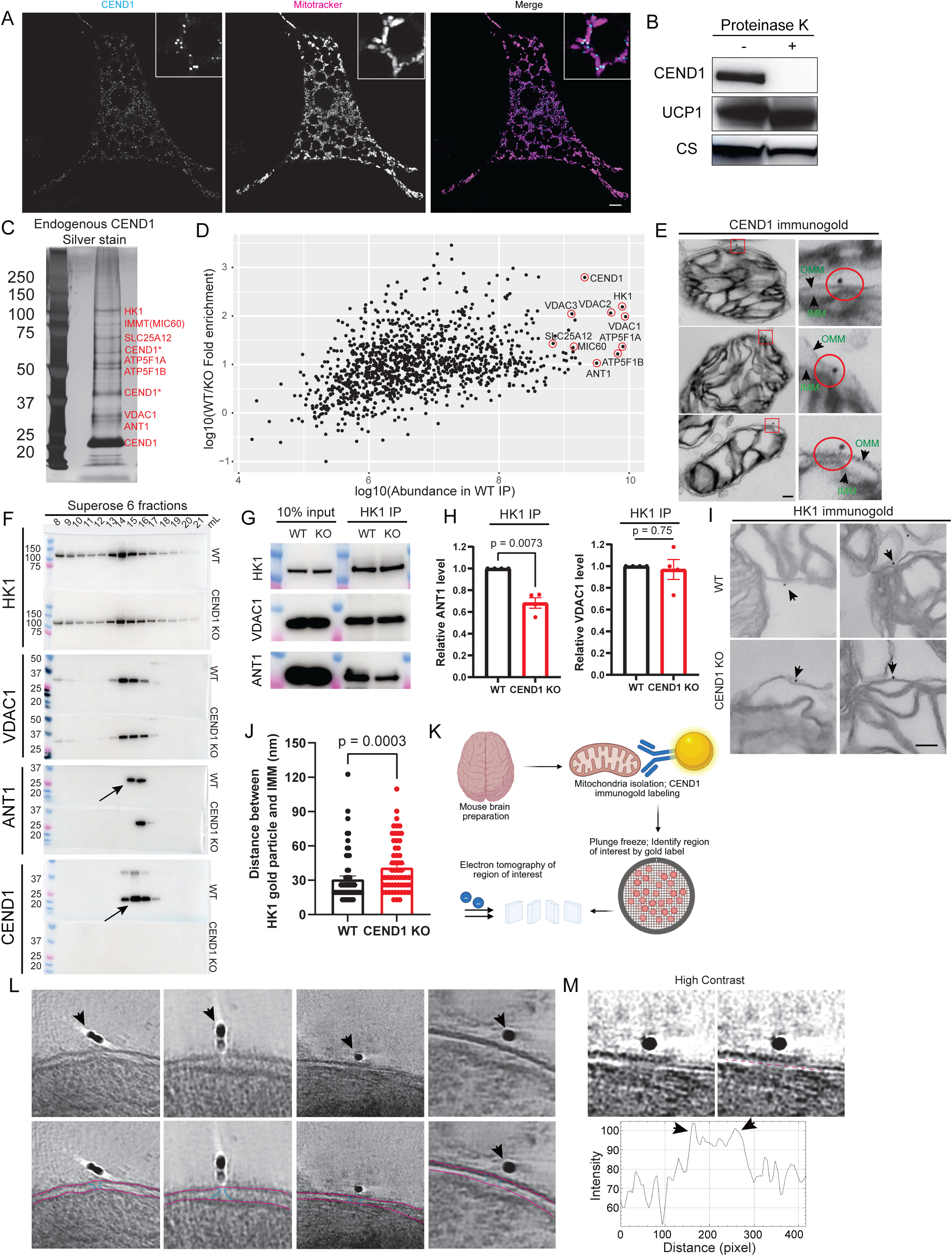
CEND1 forms a complex with HK1/VDAC1/ANT1 at OMM/IMM junctions. A. Immunofluorescence microscopy analysis of endogenous CEND1 localization in brown adipocytes. Scale bar: 5 μm. B. Proteinase K digestion of brown adipocyte mitochondria. C. Silver stain of immunopurified endogenous CEND1 complex from mouse brain mitochondrial extract. Individual bands were excised and sequenced by mass spectrometry. D. Quantitative mass spectrometry analysis of proteins isolated with the anti-CEND1 antibody from WT or CEND1 KO mouse brain mitochondrial extract. E. Immunogold electron microscopy analysis of CEND1 localization on mouse synaptosomal mitochondria. Scale bar: 100 nm. F. Gel filtration analysis of HK1/VDAC1/ANT1/CEND1 elution profiles in WT or CEND1 KO mouse brain mitochondrial extract. G-H. Western blot analysis of VDAC1 and ANT1 co-immunoprecipitated with HK1 from WT or CEND1 KO brain mitochondrial extract. A representative result is shown in G. Quantitation of 4 independent replicates are shown in H. I-J. Immunogold electron microscopy analysis of HK1 localization on mouse synaptosomal mitochondria. Scale bar: 100 nm. Representative images are shown I and quantitation is shown in J (*n* = 58 per group). K. Schematics of cryo-electron tomography analysis of mitochondrial membrane structure near CEND1. L. Representative slice images of tomograms showing density features consistent with OMM/IMM membrane bridges near CEND1. M. Line scan of the IMS showing densities consistent with OMM/IMM membrane bridges near CEND1. Data represent mean ± s.e.m. Statistical significance was calculated by two-tailed unpaired t-test (H) or Wilcoxon signed-rank test (J).

To investigate how an OMM-localized protein could influence activity-dependent glucose metabolism, we next identified potential CEND1 binding partners. Endogenous CEND1 complexes were immunopurified from mouse brain mitochondrial extracts using a monoclonal antibody against the N-terminal region of CEND1, with bound proteins competitively eluted by the antigenic peptide. SDS-PAGE followed by mass spectrometry revealed distinct bands corresponding to HK1, VDAC1, and ANT1 (Figure 5C). These proteins may together form a continuous conduit for adenine nucleotide exchange across the IMM and OMM, consistent with prior evidence in rat brain supporting the existence of such a complex^56^. IMMT (also known as MIC60), a critical regulator of cristae formation, and subunits of ATP synthase (ATP5F1A and ATP5F1B), which localize to cristae, were also detected (Figure 5C), suggesting that the CEND1/HK1/VDAC1/ANT1 complex may reside near the ATP synthesis machinery. The identification of SLC25A12 (also known as Aralar), an aspartate/glutamate antiporter and component of the MAS, points to coordination between CEND1-driven glycolysis and NADH shuttling. This is further supported by our metabolic tracing data showing higher aspartate/glutamate ^13^C-labeling in WT versus *Cend1* KO neurons (Figure 3I-J). Additionally, we observed high-molecular weight forms of CEND1, resistant to DTT reduction and detergent disruption, though their functional significance remains unclear. The specificity of the pull-down was confirmed by the marked reduction of co-purifying proteins when the same procedure was applied to mitochondria from *Cend1* KO mice (Figure 5D). Importantly, enrichment of HK1, VDAC1, and ANT1 in CEND1 pull-downs was also observed in brown adipocytes and in HEK293 cells exogenously expressing CEND1 (Figure S5D-E).

The formation of the HK1/VDAC1/ANT1 complex likely occurs at OMM/IMM contact sites, given that VDAC1 and ANT1 lack substantial domains in the intermembrane space (IMS). Notably, HK1 has been reported to localize preferentially to such contact sites where the IMS vanishes^57^. Consistent with this, immunogold electron microscopy revealed that CEND1 is frequently positioned at OMM/IMM contact sites in brain mitochondria (Figure 5E). The OMM and IMM frequently became separated, sometimes by a large distance, likely reflecting artifacts during transmission EM sample preparation process. However, CEND1-mediated formation of HK1/VDAC1/ANT1 complex may confer resistance to OMM/IMM separation, allowing the gold particles to be frequently observed at OMM/IMM contact sites.

We then analyzed how CEND1 may regulate in HK1/VDAC1/ANT1 complex formation at OMM/IMM contacts. Proteomic analysis of mitochondria from brown adipocytes or synaptosomes showed no significant changes in the abundance of proteins that co-purify with CEND1 (Figure S5F-G, supplementary table), suggesting that CEND1 deficiency does not affect their mitochondrial targeting. We thus extracted mitochondrial protein from WT or *Cend1* KO brain or BAT or from HEK293 cells with or without exogenous CEND1-expression and analyzed complex formation by gel filtration. In all systems, we observed CEND1 ablation caused a reduced amount of ANT1 co-eluting with HK1/VDAC1 (Figure 5F, S5H-J). Similarly, when we immunoprecipitated endogenous HK1 from WT and *Cend1* KO brain mitochondrial extracts, we consistently observed reduced abundance of co-purified ANT1 but not VDAC1 in the KO sample (Figure 5G-H). To demonstrate this point with an alternative approach, we performed HK1 immunogold electron microscopy on WT and *Cend1* KO brain mitochondria and quantitated the OMM/IMM distance at sites marked by the HK1 antibody. Indeed, we observed significantly larger OMM/IMM distance at HK1 sites in the KO mitochondria than the WT (Figure 5I-J), consistent with reduced HK1/ANT1 interaction. Together, these data support that CEND1 is crucial to facilitate ANT1 joining the HK1/VDAC1 complex at OMM/IMM contact sites but may be dispensable for HK1/VDAC1 interaction at the OMM.

To visualize the CEND1 complex at the OMM/IMM contact site under a more native condition at a better resolution, we performed cryo-electron tomography (cryo-ET) analysis of synaptosomal mitochondria (Figure 5K). We labeled freshly isolated mouse brain synaptosomal mitochondria with CEND1 antibody and gold-labeled protein A to aid the localization of CEND1 on the OMM and collected tilted series. Examining the reconstructed tomograms revealed density connecting the OMM and IMM where CEND1 was found to localize (Figure 5L). A line scan along the IMS revealed extra intensity along the edge of the density (Figure 5M), suggesting hemifusion-like junction formation at OMM/IMM contact sites. These observations provide further evidence supporting the presence of the CEND1 complex at the OMM/IMM interface.

### CEND1 couples mitochondrial ATP production and HK1 ATP consumption

We next investigated the functional significance of CEND1 in promoting HK1/VDAC1/ANT1 complex formation. Our hypothesis is that this complex provides a passageway enabling efficient ATP/ADP exchange between mitochondrial matrix and OMM-bound HK1, thereby ensuring targeted and efficient ATP delivery to HK1 in neurons and brown adipocytes under stimulation.

We first tested this in brown adipocyte mitochondria. Unlike most cell types, brown adipocyte mitochondria generate a substantial fraction of ATP through substrate-level phosphorylation at the succinyl-CoA synthetase step, with an apparent P/O ratio of ∼0.2 when respiring on pyruvate/malate^9^. This is attributable to the predominant expression of the ATP-forming isoform of succinyl-CoA synthetase (SUCLA2) in brown adipocytes (Figure S6A, supplementary table). To evaluate how CEND1 affects HK1 activity using intramitochondrially derived ATP, we isolated mitochondria from WT and *Cend1* KO BAT and supplied them with glucose, ADP, and either α-ketoglutarate or succinate while measuring glucose-6-phosphate (G6P) production. Although both substrates supported equally robust respiration (Figure S6B), only α-ketoglutarate could support sustained ATP production through substrate-level phosphorylation because it feeds into the SUCLA2-catalyzed step (Figure S6C). Consistent with this, α-ketoglutarate supported significantly higher G6P production than succinate, and the rate was higher in WT than KO mitochondria (Figure 6A). Importantly, ATP production itself was comparable between WT and KO mitochondria supplied with α-ketoglutarate (Figure S6D), indicating that reduced HK1 activity in the KO is due to inefficient ATP delivery rather than impaired production.

**Figure 6.**
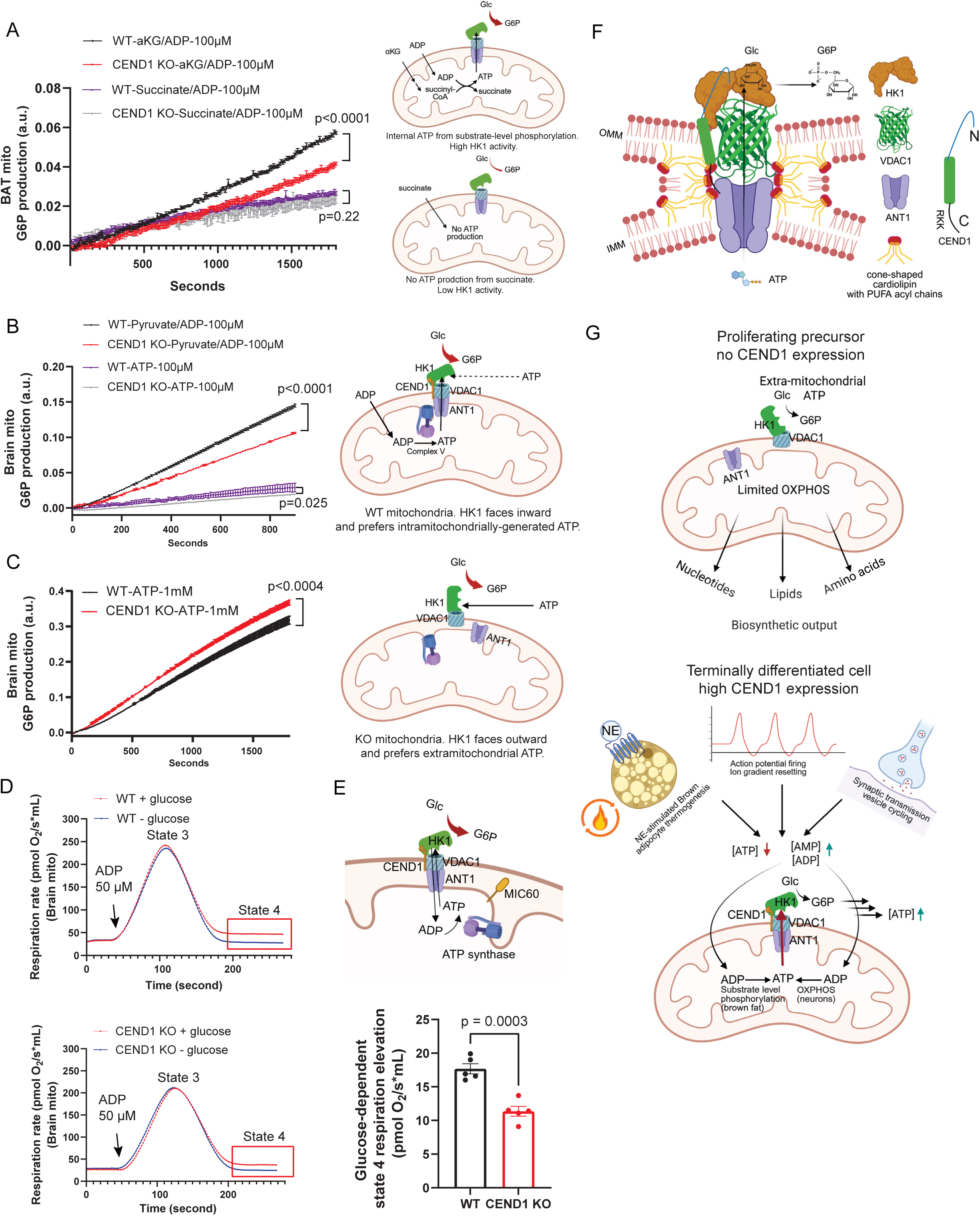
CEND1 couples mitochondrial ATP production and HK1 ATP consumption. A. Brown adipocyte mitochondrial HK1 activity assay in the presence of 10 mM α-ketoglutarate or succinate and 100 μM ADP (*n* = 5 per group). The schematics on the right show how α-ketoglutarate but not succinate supports ATP production via substrate-level phosphorylation. B. Brain mitochondrial HK1 activity assay in the presence of 10 mM pyruvate/5 mM malate/ and 100 μM ADP or 100 μM ATP (*n* = 5 per group). The schematics on the right show how CEND1 promotes HK1 utilization of intramitochondrially-derived ATP. C. Brain mitochondrial HK1 activity assay in the presence of 1 mM ATP (*n* = 5 per group). The schematics on the right show how the absence of CEND1 enhances the accessibility of extra-mitochondrial ATP to HK1. D. Representative oxygen consumption traces of WT and CEND1 KO brain mitochondria +/- 10 mM glucose. Note glucose addition caused a persistently higher State 4 respiration rates. E. Schematic illustration and quantitation of glucose-induced elevated State 4 respiration rates (*n* = 5 per group). Data are mean ± s.e.m. Statistical significance was calculated by two-way ANOVA with repeated measurements (A, B, C, and D) or unpaired t-test (E). Model of the HK1/VDAC1/ANT1/CEND1 complex housed in a hemifusion-like structure at OMM/IMM contact site. G. Model of how CEND1 directs HK1 ATP preference to facilitate activity-dependent glucose metabolism as mitochondria shift from a biosynthetic role in proliferating precursors to the terminal differentiated state when response to stimulation becomes necessary.

We next examined brain mitochondria, where oxidative phosphorylation is the main mechanism of ATP production. WT mitochondria respiring on pyruvate/malate with 100 μM ADP displayed robust G6P formation (Figure 6B). Strikingly, the addition of 100 μM ATP instead of ADP dramatically reduced HK1 activity (Figure 6B), consistent with HK1’s preference for ATP delivered internally via VDACs. Compared with WT, *Cend1* KO mitochondria displayed significantly lower HK1 activity with ADP as the substrate (Figure 6B), despite producing similar amounts of ATP (Figure S6E). Thus, as in brown adipocytes, the deficit in the KO reflects impaired ATP delivery to HK1. Interestingly, at high concentrations of external ATP (1 mM), KO mitochondria showed higher G6P production than WT (Figure 6C). Since CEND1 deficiency does not impair HK1’s binding to VDAC1 or its localization to the OMM, this observation is consistent with a scenario in which CEND1 promotes an HK1 conformation that orients its ATP-binding site toward VDAC1, enforcing utilization of matrix-derived ATP while limiting access to cytosolic ATP. This model also provides a mechanistic explanation for the trend towards higher ^13^C-pyruvate and lactate labeling and ATP levels observed in KO neurons under resting conditions (Figure S3C-D, Figure 4A).

Our mitochondrial HK1 activity assays support the idea that the CEND1/HK1/VDAC1/ANT1 complex establishes a closed ATP/ADP cycling loop. A key prediction is that this continuous recycling should sustain mitochondrial respiration under state 4 conditions. To test this, we performed high-resolution respirometry on WT and KO brain mitochondria in the presence or absence of glucose. In the absence of glucose, both WT and KO mitochondria displayed a transient increase in respiration upon ADP addition (state 3), which declined rapidly to baseline due to ADP depletion (state 4) (Figure 6D). In contrast, when glucose was supplied, both WT and KO mitochondria exhibited markedly elevated state 4 respiration, consistent with persistent ADP regeneration by the HK1 reaction (Figure 6D-E). Notably, the magnitude of this glucose-induced state 4 elevation was significantly greater in WT mitochondria, indicating more efficient ATP/ADP cycling through the VDAC1/ANT1 conduit rather than dissipation into the surrounding medium (Figure 6D-E).

Together, these results demonstrate that CEND1 couples mitochondrial ATP production to HK1 activity in both neurons and brown adipocytes. While the two cell types differ in their ATP production strategies, substrate-level phosphorylation in brown adipocytes versus oxidative phosphorylation in neurons, CEND1 enables efficient and localized ATP channeling to HK1 across both contexts.

## Discussion

Our study identifies CEND1 as a previously unrecognized regulator of activity-dependent glucose metabolism in both brown adipocytes and neurons. We show that CEND1 is required for rapid, stimulation-evoked ATP production in brown adipocytes, supporting cAMP synthesis and glycolytic upregulation during adrenergic activation. Remarkably, CEND1 performs an analogous function in neurons: its loss blunts activity-dependent glucose utilization, diminishes ATP regeneration, and reduces neuronal firing, ultimately raising the seizure threshold. Mechanistically, CEND1 forms a complex with HK1, VDAC1, and ANT1 at OMM/IMM junctions, enabling efficient coupling of mitochondrial ATP production to HK1-mediated glucose phosphorylation (Figure 6F). Together, these findings reveal CEND1 as a conserved metabolic scaffold that links mitochondrial ATP output to glucose utilization in highly active, stimulation-responsive, and energy-demanding cells.

Our cryo-ET analyses suggest that the CEND1 complex resides within a hemifusion-like membrane bridge formed at OMM/IMM contact sites (Figure 6F). This interpretation is reinforced by observations from other systems, including frequent membrane plane skipping in freeze-fracture EM^58^ and selective enrichment of PUFA-acylated cardiolipins in OMM/IMM contact-site fractions^59^. PUFA-acylated cardiolipins possess a pronounced cone-shaped geometry, stemming from their small headgroup and the large cross-sectional area of highly flexible acyl chains, which is generally incompatible with planar bilayers but ideally suited to support the strong negative curvature required for hemifusion-like membrane bridges^60^. Such curvature-stabilizing properties align well with the proposed architecture that accommodates the CEND1 complex. Notably, the C-terminal polybasic RKK motif of CEND1 is well positioned to engage with cardiolipin’s negatively charged headgroups, potentially stabilizing such hemifusion-like structures (Figure 6F), analogous to our prior findings that the R-rich region of CLSTN3B stabilizes hemifusion-like ER-lipid droplet membrane bridges^61^. Formation of a CEND1-supported hemifusion architecture would be advantageous for directed ATP channeling by stabilizing the VDAC1/ANT1 coupling and constraining local ATP diffusion. Additionally, brain mitochondrial cardiolipin is enriched in arachidonic acid (20:4) and docosahexaenoic acid (DHA, 22:6)^62^, and epidemiological data link high dietary DHA intake, such as from deep-sea fish, to reduced Alzheimer’s disease risk^63–66^. Whether PUFA-induced increases in membrane fluidity and enhanced hemifusion architecture at OMM/IMM contact-site contribute to improved neuronal metabolic responsiveness and cognitive function remains an intriguing possibility that warrants further study.

The elevated seizure threshold in CEND1-deficient mice supports a functional link between neuronal glucose metabolism and seizure susceptibility. The anti-epileptic benefits of the fasting or ketogenic diet have long been recognized^67^. Whereas the mechanisms underlying the ketogenic diet-mediated seizure suppression are multifaceted^68–73^, our understanding of CEND1 function suggests that glucose availability may be a critical component. Because CEND1 supports activity-dependent ATP production from glucose, the reduced excitability and increased seizure resistance in *Cend1* KO neurons may stem from impaired glucose-driven ATP generation during neuronal firing. This ATP deficit could dampen action potential propagation or promote opening of K_ATP_ channels, both of which reduce neuronal excitability. On the other hand, CEND1-mediated activity-dependent glucose-derived carbon fluxes may enhance neuronal excitability by supporting neurotransmitter replenishment, particularly glutamate, as indicated by CEND1-dependent ^13^C labeling of glutamate (Figure 3J). Future neurophysiological studies directly comparing neuronal activity in glucose versus ketone body conditions will help determine whether the defects observed in *Cend1* KO neurons reflect a specific inability to utilize glucose as an energy source during activation.

Our findings also provide insight into to the long-standing debate regarding whether neurons primarily rely on glucose or astrocyte-derived lactate to sustain activity. The classical view posits direct neuronal glucose oxidation, whereas the astrocyte-neuron lactate shuttle (ANLS) model proposes preferential neuronal use of lactate released from activity-stimulated astrocytes^6,74,75^. The neuronal selective expression of CEND1, however, indicates that intrinsic neuronal glucose metabolism plays a prominent role in supporting rapid, activity-dependent ATP production. Notably, arguments against significant neuronal glycolysis have focused on low neuronal PFKFB3 expression^76^, which limits flux through PFK1. Yet neuronal glucose catabolism can proceed through the pentose phosphate shunt (PPP), which bypasses PFK1 and re-enters glycolysis at the glyceraldehyde-3-phosphate step. This route simultaneously supports energy generation and provides NADPH for redox homeostasis, a necessity given the vulnerability of neurons to oxidative stress. Indeed, enforced PFKFB3 expression enhances oxidative damage, highlighting the importance of this alternative metabolic routing^77^. Note that our CEND1 model does not exclude neuronal lactate utilization. Rather, astrocyte-derived lactate could provide an immediate substrate to sustain mitochondrial respiration and oxidative phosphorylation during early stimulation to jump start the ATP supply to neuronal HK1, in which case neuronal glucose metabolism and astrocytic lactate provision may operate cooperatively and in a temporally coordinated manner to meet dynamic energetic demands.

The enhanced access of HK1 to extra-mitochondrial ATP in the absence of CEND1 highlights dual functionality: CEND1 not only promotes activity-dependent glucose metabolism by coupling HK1 to mitochondrial ATP production but also restricts resting-state glucose utilization by restricting HK1 access to cytosolic ATP when mitochondrial respiration is low. This scenario is further corroborated by elevated resting state ^13^C labeling of pyruvate/lactate and ATP levels in *Cend1* KO neurons. The dichotomous effect of CEND1 on glucose metabolism appears to be tuned by its unique transcriptional regulation, allowing cells to adapt to specific metabolic demands. For example, in brown adipocytes, although CEND1 is induced during cold exposure, it is paradoxically strongly repressed by LSD1. We speculate that in resting brown adipocytes, LSD1-mediated repression of CEND1 may preserve glycolytic flux when mitochondrial activity is primarily directed toward *de novo* lipogenesis and LD expansion rather than ATP generation via the SUCLA2 reaction to support HK1 reaction. Consistent with this, we observed markedly increased lipid accumulation upon CEND1 ablation in LSD1-deficient brown adipocytes, suggesting that enhanced cytosolic ATP availability to HK1 drives greater flux of glucose-derived carbon into lipids in the absence of CEND1. LSD1-mediated repression of CEND1 was also observed in developing cardiomyocytes and de-repression of CEND1 following LSD1 ablation impairs cardiomyocyte precursor proliferation^78^, reminiscent of studies showing that CEND1 promotes cell-cycle exit in neural progenitors^34^. These findings suggest that CEND1-mediated coupling of HK1 to mitochondrial ATP production may be optimized for terminally differentiated, stimulus-responsive cells but can be detrimental in proliferating precursors. By forcing HK1 to rely on mitochondrial rather than cytosolic ATP, CEND1 may constrain glycolytic and biosynthetic processes in proliferating cells, which derive a large part of ATP from glycolysis while depending on anabolic mitochondrial function for synthesizing cellular building blocks (Figure 6G)^79^. This constraint likely underlies the anti-proliferative effects of CEND1 observed in cardiomyocyte and neural progenitors. Interestingly, LSD1 is essential for neuronal progenitor proliferation^80^, raising the possibility that repression of CEND1 may also contribute to this process. Overall, epigenetic regulation of CEND1 enables cells to coordinate HK1-driven glucose metabolism with mitochondrial operational modes (catabolic vs anabolic) and ATP-producing capacity thereby allowing adaptation to their differentiation and activity states.

## Methods

### Mouse strains

The *Cend1^-/-^* and the adipose-specific *Lsd1* KO mice were described in previous publication (Sergaki et al, PMID: 20153830; Zeng et al, PMID: 31043739). Mice were maintained on a normal chow diet (13% fat, 57% carbohydrate and 30% protein, Teklad Rodent Diet 5L0D). All mice were maintained under a 12 hr light/12 hr dark cycle at constant temperature (23°C or 30°C as specified) and 40–60% humidity with free access to food and water. All animal studies were approved by and in full compliance with the ethical regulation of the Institutional Animal Care and Use Committee (IACUC) of University of Texas Southwestern Medical Center. Mice were euthanized by CO_2_ asphyxiation, according to the guidelines of IACUC and the recommendations of the Panel on Euthanasia of the American Veterinary Association. Sample size was chosen based on literature and pilot experiment results to ensure that statistical significance could be reached. Randomization was not performed because mice were grouped based on genotype. Littermates were used for all the experiments involving the conditional KO mice.

### Antibody

HK1 (Proteintech, 19662-1-AP); HK1 (Invitrogen, MA5-14789); HK1 (Abclonal, A0533); VDAC1 (Proteintech, 55259-1-AP); ANT1/SLC25A4 (Cell Signaling, 51755); SLC25A4 (Proteintech, 30631-1-AP); CEND1 (Cell signaling, 8944); CoraLite® Plus 488-conjugated TOM20 Polyclonal antibody (Proteintech, CL488-11802); Neuron-specific beta -III Tubulin Antibody (RD systems, MAB1195); UCP1(Abcam, ab209483); CS (Abcam, ab129095). FLAG (Sigma, F1804), Alexa Fluor 488 Fluor goat anti-Rabbit (Invitrogen, A11034), Alexa Fluor 488 Fluor goat anti-mouse (Invitrogen, A-11001); Peroxidase AffiniPure® Goat Anti-Rabbit IgG (H+L) (Jackson ImmunoResearch, 111-035-003); Peroxidase AffiniPure® Goat Anti-Mouse IgG (H+L) (Jackson ImmunoResearch, 115-035-003)

### Fixed cell imaging

Primary brown adipocytes, neurons and Hela cells plated onto glass coverslips were stained with Mito tracker dye (Invitrogen, M7512) and used for fixed cell imaging. Images were taken on a ZEISS 900 confocal microscope with Airyscan.

### Cold tolerance test

Mice housed at 3 to 5 per cage were pre-acclimatized at thermoneutrality (28-30□°C) for two weeks before being shifted to 4□°C. During cold tolerance test, mice were singly housed without access to food. Hourly body temperature was recorded using a microprobe thermometer (Physitemp, BAT-12) equipped with a rectal probe (Physitemp, RET3).

### NE turnover in BAT

Mice were i.p. injected with 120 mg/kg of TH inhibitor α-methyl-p-tyrosine-methyl ester (Sigma M3281) immediately prior to being transferred into a 4°C incubator. At indicated time points, mice were sacrificed and BAT dissected and homogenized in 500 μL 1 M HClO_4_ per lobe. The homogenate was neutralized with 2 M KOH/0.2 M MOPS and centrifuged to remove KClO_4_ precipitants. NE content in the supernatant was analyzed by HPLC (Agilent 1260 Infinity II) coupled with an electrochemical detector/SenCell flow cell (Antec Scientific DECADE Elite). Mobile phase:100 mM acetic acid, 20 mM citric acid, 0.2 mM EDTA, 7.5% MeOH, 2 mM NaCl, 500 mg/L octanesulfonic acid sodium salt, pH to 4.86 with NaOH. Working potential of the flow cell was set at 0.5V.

### Primary brown adipocyte assays

Primary adipocytes were isolated as previously described (Zeng et al, PMID: 31043739) and used as suspension for activity assays or attached to glass coverslips pre-coated with laminin and poly-D-lysine for fixed cell imaging.

Respiratory responses to norepinephrine was measured by high-resolution respirometry with an Oroboros Oxygraph-2k (Oroboros Instruments, Innsbruck, Austria) in a standard configuration, with 2 ml volume of the two chambers, at 37°C, and 350 rpm stirrer speed. Specifically, freshly isolated primary brown adipocytes were suspended at a density of 1-2 ×10^6^ cells/mL in Krebs-Ringer modified buffer (KRMB) consisting of 120 mM sodium chloride, 5 mM potassium chloride, 1 mM magnesium chloride, 1.3 mM calcium chloride, 0.4 mM dibasic potassium phosphate, 20 mM HEPES, 10 mM sodium bicarbonate, 10 mM glucose and 4% (w/v) BSA (fatty acid free) at pH 7.3. For each measurement, 100,000 cells suspending in 2 mL of KRMB were added to the respiration chamber. Basal respiration rate was determined first, followed by injecting 1 μM NE into the chamber to stimulate respiration. The software DatLab7 was used for data acquisition (2s time intervals) and analysis. All measurements were performed within 2 hours of cell isolation.

For cAMP quantitation, primary brown adipocytes were suspended in 300 µL KRMB at 10^6^ cells/ml density ± 1μM NE. Three replicate measurements were set up per condition. After 5 minutes at 37°C on a shaker, the brown adipocyte suspension was transferred to a centrifugal filter unit (EMD Millipore UFC30HV00) and immediately centrifuged at 10,000 rpm for 30 s to separate the cells and the buffer. Following the centrifugation, the adipocytes collected in the filter unit were immediately lysed with 300 μL 0.1 M HCl. The cAMP levels of brown adipocytes were then measured using a Cyclic AMP ELISA kit (Cayman Chemical, 581001) following manufacturer’s protocol. To inhibit cAMP hydrolysis, 250 μM 3-isobutyl-1-methylxanthine (IBMX) (Cayman, 13347) was included in NE samples.

For lactate measurement, the brown adipocyte suspensions at indicated time points were transferred to a centrifugal filter unit (EMD Millipore UFC30HV00) and immediately centrifuged at 10,000 rpm for 30 s to separate the cells and the buffer. The lactate levels in buffer were measured by a Lactate-Glo Assay kit (Promega, J5022) following manufacturer’s protocol.

### Nucleotide quantitation

Primary brown adipocytes suspended in 500 µL KRMB at 10^6^ cells/ml density ± 1μM NE were incubated at 37°C on a shaker. After 5 minutes, the brown adipocyte suspension was transferred to a centrifugal filter unit (EMD Millipore UFC30HV00) and immediately centrifuged at 10,000 rpm for 30 s to separate the cells and the buffer. Following the centrifugation, cells were extracted with 500 µl solvent containing Methanol:Acetonitrile:Water 2:2:1 (v/v/v). The extract was transferred to a new 1.5ml Eppendorf tube and subjected to three freeze-thaw cycles between liquid nitrogen and 37°C water bath. After the third thaw, the samples were vortexed for 1 min and then centrifuged at a maximum speed for 15 minutes in a refrigerated centrifuge. The supernatant was transferred to a new tube and dried in a SpeedVac to a volume just below 100 μL. The ATP levels were then measured using the CellTiter-Glo Luminescent Cell Viability Assay kit (Promega, G7570) following manufacturer’s protocol. To measure the ADP and AMP levels, the ADP in samples was converted into ATP by the addition of phosphoenolpyruvate (PEP, Sigma, P7127) and pyruvate kinase (Sigma, P9136), and the AMP was further converted into ATP by the addition of adenylate kinase (Sigma, M3003). And these conversions were validated by anion-exchange chromatography approach. For this approach, the brown adipocyte and adenine nucleotide standard samples were diluted to 500 μL by adding 5 mM Tris-HCL pH 8.0 and resolved on Capto HiRes Q 5/50 column 0-40%B in 15 mL (Buffer A: 5 mM Tris-HCl 8.0; Buffer B: 5mM Tris-HCl pH 8.0 and 1M NaCl). The ATP, ADP and AMP peaks were monitored and quantified based on UV absorption at 254 nm.

Primary cortical neurons were pre-acclimated to Tyrode’s buffer (pH7.4 at 37oC) for 10 minutes before electrical stimulation. Electrical stimulation was performed at 10 Hz, 1 min duration, 3 minutes later, samples were collected in 500 μL 80% Methanol. Adenine nucleotides were extracted, processed and measured by the same approach as used in brown adipocytes. IMP levels were determined by a metabolomics approach.

### PET-MRI analysis of ^18^F-FDG uptake in BAT

Each mouse received 0.25 mCi of [^18^F]-FDG (volume 130 µL) and 1 mg/kg CL316,243 via lateral tail-vein injection. After injection, mice remained awake for a 20-min uptake period in a warmed cage (30°C). All imaging was performed on a Bruker BioSpec 7.0-T MRI system integrated with a Bruker PET insert Si 103 (axial FOV 101 mm, 3 Rings). A mouse quadrature volume RF coil (35-mm ID) was used for all MR acquisitions. PET and MRI were performed simultaneously with the mouse positioned prone on a heated animal bed equipped with continuous isoflurane delivery.

Mice were anesthetized with 1.5-2.0% isoflurane in medical air during imaging. Respiratory rate and body temperature were continuously monitored using a small-animal monitoring system (SAII, Stony Brook, NY). Body temperature was maintained at 36-37°C using a feedback-controlled warm-air system.

For MRI Acquisition, an initial localizer scan was followed by the following sequences: T1 weighted 3D FISP (TR/TE = 5.5/2.6 ms, Number of Segments = 210, FOV = 40 x 40 x 75 mm, Matrix = 256 x 256 x 64); T2-weighted Turbo RARE (TR/TE = 1000/27 ms, RARE factor = FOV = 40 x 40 mm, Matrix = 256 x 256, Slice Thickness = 1mm). MRI datasets were used for anatomical localization, PET attenuation correction, and region-of-interest (ROI) definition.

For PET Acquisition, simultaneous PET data were acquired for 30 minutes beginning 45 minutes post-injection. PET data were reconstructed using an OSEM-3D algorithm with corrections for detector normalization and decay.

PET and MRI datasets were co-registered and analyzed using PMOD image processing software (PMOD Technologies LLC, Zurich, Switzerland; version 4.403). Brown adipose tissue was identified on co-registered T2 weighted MRI images and ROIs were manually drawn to calculate Standardized Uptake Values (SUVs).

### UP state recordings and analysis

UP state experiments were performed in acute somatosensory neocortical slices prepared from male (P18-24) littermates of each genotype. Slice preparation was based on our earlier study (Hays et al, PMID: 21976507). Male WT and Cend1 KO littermates were deeply anesthetized with ketamine/xylazine cocktail (4 mg/ml xylazine in 30 mg/ml ketamine) and decapitated. The brain was transferred into an ice-cold dissection buffer containing (in mM): 87 NaCl, 3 KCl, 1.25 NaH_2_PO_4_, 26 NaHCO_3_, 7 MgCl_2_, 0.5 CaCl_2_, 20□D-glucose, 75 sucrose, and 1.3 ascorbic acid. The buffer was constantly aerated with 95% O_2_-5% CO_2_. The brain was glued to an angled block (Agmon and Connors, PMID: 1870696). Thalamocortical slices, 400 μm, were cut using a vibratome (Leica VT 1200S). Slices were immediately transferred to an interface recording chamber (Harvard Instruments) and allowed to recover for 1 h in nominal artificial CSF (ACSF) at 32°C containing (in mM): 126 NaCl, 3 KCl, 1.25 NaH_2_PO_4_, 26 NaHCO_3_, 2 MgCl_2_, 2 CaCl_2_, and 25□D-glucose. Data acquisition was controlled using custom Labview software available at https://github.com/jaygibson244/GibsonHuber-UP-states.git.

After the recovery described above, slices were perfused with a modified ACSF that better mimics physiological ionic concentrations□in vivo 3 containing (in mM): 126 NaCl, 5 KCl, 1.25 NaH_2_PO4, 26 NaHCO_3_, 1 MgCl_2_, 1 CaCl_2_, and 25□D-glucose (based on but modified from□Hays et al, PMID: 21976507 and Sanchez-Vives et al, PMID: 11017176). Slices remained in this modified ACSF for 45 min followed by the UP state recordings.

Spontaneously generated UP states in brain slices□were extracellularly recorded using 0.5 MΩ tungsten microelectrodes (FHC) placed in layer 4 of primary somatosensory cortex. Only 1 recording was obtained from each slice and data was collected from 4-8 slices per mouse. This extracellular monitoring of UP states is a reliable indicator of the synchronous, depolarized state of neuron populations from which the term “UP state” was originally defined (Sanchez-Vives et al, PMID: 11017176; Rigas et al, PMID: 26528142). A total of 5 min of spontaneous activity was collected from each slice. Recordings were amplified 10,000-fold, sampled at 2.5 kHz, and filtered on-line between 500 Hz and 3 kHz. All measurements were analyzed off-line using custom Labview software. This code is available at https://github.com/jaygibson244/GibsonHuber-UP-states.git. For visualization and analysis of UP states, traces were offset to zero, rectified, and low-pass filtered with a 0.2 Hz cutoff frequency. Using these processed traces, the threshold for detection was set at 15× the RMS (root mean square) noise. An event was defined as an UP state if its amplitude remained above the threshold for at least 200 ms. The end of the UP state was determined when the amplitude decreased below threshold for >600 ms. Two events occurring within 600 ms of one another were considered a single UP state. Slices were excluded if <2 UP states occurred per minute. UP state amplitude was defined based on the filtered/rectified traces. This amplitude may be considered a coarse indicator of the underlying firing rates of neuronal populations. Normalized power for 3 frequency bands - 30-55, 65-100, and 200-600 Hz - was determined by calculating the power function over time only during an UP state and then normalizing this function by total power in the 13-600 Hz window. Direct measures of firing rates were not possible because individual spikes could not be isolated except during the quiet periods (the DOWN states).

### Simultaneous fEPSP and glucose sensor recordings and analysis

Acute hippocampal brain slices were prepared from P30-35 wild-type (WT) and Cend1 KO littermates as described previously (Jakkamsetti et al, PMID: 24094104). Briefly, mice were anesthetized with ketamine (120 mg/kg)/xylazine (16 mg/kg) and transcardially perfused with chilled (4 °C) sucrose dissection buffer (as for the acute cortical slices preparation). Transverse hippocampal slices (400 µm) were obtained on a Leica VT1200S slicer. Slices were recovered for at least 2 hours in a submerged recovery chamber and maintained at 30 °C in artificial cerebral spinal fluid (ACSF) containing the following (in mM): 126 NaCl, 5 KCl, 1.25 NaH_2_PO_4_, 26 NaHCO_3_, 1 MgSO_4_, 2 CaCl_2_, and 5□D-glucose aerated with 95% O_2_/5% CO_2_ to pH 7.4.

For all recordings, slices were submerged and perfused with ACSF at 2.5–3.5 mL/min (30□±□1 °C). Field potentials (fEPSP) were evoked by stimulation of the Schaffer collateral pathway with a concentric bipolar electrode (FHC; catalog # 30205) and recorded with a glass electrode (1 MΩ) filled with ACSF, placed in the stratum radiatum of CA1. Test stimuli were delivered every 30 s. The glucose sensor probe (Pinnacle, 25 μm diameter glucose micro-biosensor, 7020-L-25-Glucose) was placed within 100 µm of the recording electrode. fEPSP recordings were performed at the same sampling rate and filter settings described above. Extracellular glucose was sampled at 1 Hz. Each stimulation was a bipolar pulse (100 ms up, 100 ms down). A single train of 150 of these pulses were applied at a rate of 10 Hz (total of 15 seconds). The fEPSPs were recorded for those 15 seconds. The glucose levels were recorded for 7 minutes after stimulation onset. This was repeated 6 times in a single experiment, but with increasing stimulation intensities (in μA: 2,4,8,15,30,60). As expected, responses to each stimulation pulse had a negative polarity (Angamo et al, PMID: 27559140). Both fEPSPs and popspikes were measured when both existed – with the latter being a second, longer latency negative hump in the response waveform (Angamo et al, PMID: 27559140).

### Seizure threshold measurement

Mice were transferred to the laboratory for 1 h before the experiment to habituate to the environment. All flurothyl tests were performed between 0900 and 1200-h in a ventilated chemical fume hood to minimize circadian rhythm influences on flurothyl-induced seizure thresholds. Individual mouse was placed in an air-tight acrylic cylinder chamber (Ted Pella, model: 2240-1) for 1 min before inducement. Generalized tonic-clonic seizures were elicited using a 10% flurothyl (Sigma-Aldrich, 333-36-8) dissolved in 95% ethanol. Flurothyl was infused at a rate of 200 µL/min on to a disk of filter paper 3 (Whatman, Grade1) suspended at the top of the chamber using a 20 mL glass syringe (Sigma-Aldrich, Z101079) driven by an infusion pump (Sage Instruments, White Plains, NY). Each test was video-recorded and lasted for up to 10 minutes or until the animal begins to exhibit tonic-clonic seizure activity (sustained loss of posture control for greater than 2 s accompanied by the trunk rigidity). Immediately upon completion of data acquisition, infusion was terminated and the lid of the chamber was removed, exposing the mouse to room air. On each trial, the latency (in seconds) from the start of the infusion to the loss of postural control was used to indicate the generalized seizure threshold.

### Primary neuron isolation

Primary mouse cortical neurons were prepared using a Pierce™ Primary Neuron Isolation Kit (Thermo Scientific™, 88280) following manufacturer’s protocol with some modifications. Briefly, cortices of P0 mouse pups were micro-dissected, papain-digested for 30 minutes at 37°C in an incubator and triturated in low ovomucoid solutions to get single cell suspensions. Then high ovomucoid solution was added slowly to the bottom of triturated cell solution. Cells were then centrifuged down to the high ovomucoid solution at 1000 rpm for 15 minutes at room temperature, resuspended in pre-warmed Serum-supplemented Neuronal Culture Medium (10% FBS, 2 mM L-Glutamine and 1% Pen/Strep), and seeded on glass coverslips placed in a 12-well plate (Neuvitro, GG181.5LAMININ). After 24 hours, replace the Serum-supplemented Neuronal Culture Medium with an equivalent volume of Serum-free Neuronal Culture Medium (B27 supplement, 2 mM L-Glutamine and 1% Pen/Strep). At day 3, Neuronal Growth Supplement was added to each well. Cultures were re-fed every three days by replacing half of the media. In experiments requiring fixed cells, cortical neurons were fixed on DIV11 with 4% PFA, stored in cold PBS for immunofluorescence. In experiments for western blot, cortical neurons on DIV 5, 8 and 11 were collected from the coverslip using a RIPA buffer.

### ^13^C glucose isotope tracing

For ^13^C-U-glucose (Cambridge Isotope Labotatories, CLM-1396-1) tracing experiment, primary cortical neurons cultured on DIV 11 were pre-acclimated to Tyrode’s buffer (pH7.4 at 37°C) containing 119 mM NaCl, 2.5 mM KCl, 10 mM glucose, 50 mM HEPES, 2 mM CaCl_2_, 2 mM MgCl_2_ for 10 minutes. Then the neurons were switched to the Tyrode’s buffer containing 10mM U-^13^C-glucose with or without receiving an electrical stimulation for 1 minute (10 Hz). After nine minutes, neurons were harvested with 500 µL ice cold 80% methanol. Metabolites were extracted with three freeze-thaw cycles in liquid nitrogen. Supernatant was collected after a 15 min centrifugation at 13,000g at 4 °C and lyophilized. Metabolites were derivatized with trimethylsilyl (TMS) at 42 °C for 30 min for GC-MS analysis. ^13^C-enrichment analysis was performed by GC-MS as previously described (Mullen et al, PMID: 24857658).

### CEND1 or HK1 complex purification

Mitochondria isolated from brain, BAT or HEK 293 cells transfected with FLAG-CEND1 were extracted with 1% or 1.5% n-dodecyl-β-D-maltoside (DDM, Anatrace, D310) lysis buffer containing 20 mM Tris-HCl pH8.0 and 150mM NaCl supplemented with protease Inhibitors for 1 h at 4°C with constant rotating. The cleared lysate by centrifugation at 12,000g for 5 min was incubated with Protein G dynabeads (Invitrogen, 10004D) pre-conjugated with CEND1 antibody (Cell signaling, 8944) or anti-FLAG-M2 magnetic beads (Millipore, M8823) for 3 hr at 4°C with constant rotating for immunoprecipitation. After incubation, the dynabeads or resin were washed 3 times with 0.1% DDM wash buffer containing 20 mM Tris-HCl pH8.0 and 150 mM NaCl, the CEND1 complex was then eluted for 20 min twice with 200 µL 100 µg/ml CEND1 antigen peptide (Peptide Synthesis by ThermoFisher Scientific) or 3x FLAG peptide (Millipore, F3290) in 0.1% DDM wash buffer. Elutes were collected and concentrated to around 20 µL by using a 50 kDa Amicon® Ultra Centrifugal Filter (Millipore, UFC5050) for SDS gel silver staining by using a commercial kit (Thermo Scientific, 24612) or proteomics analysis.

For HK1 complex isolation, the cleared lysate by centrifugation was incubated with Protein G dynabeads (Invitrogen, 10004D) pre-conjugated with HK1 antibody (Abclonal, A0533) for 3 hr at 4oC with constant rotating for immunoprecipitation. After incubation, the dynabeads were washed 3 times with 0.1% DDM wash buffer containing 20 mM Tris-HCL pH8.0 and 150mM NaCl, then the beads were eluted by 100 µl TEAB buffer with 5% SDS for 10 min at 70oC. Elutes were collected and used for Western blotting.

### Mitochondrial isolation

All mitochondria isolation and fractionation were carried out on ice. Crude brain mitochondria were isolated from adult mouse forebrain by homogenization in an isolation buffer (225 mM sucrose, 75 mM mannitol, 1 mM EGTA and 5 mM HEPES pH 7.4) and centrifugation and were then purified by Percoll discontinuous gradient centrifugation as previously described (Kristian et al, PMID: 20578034). Isolated mitochondrial pellet was used for protein extraction or resuspended in a respiration buffer (RES) containing 100 mM KCl, 2 mM MgCl_2_, 1 mM EDTA, 4 mM K_2_HPO_4_, 0.1% BSA-fatty acid free, 20 mM HEPES pH 7.2 for mitochondrial HK1 activity assay or respiration assay.

Mouse BAT were dissected, minced, and further homogenized with a motorized Teflon pestle in ice-cold HES buffer (20 mM HEPES, 1mM EDTA, 250 mM sucrose, pH 7.2) supplemented with 1% fatty acid-free BSA (Millipore, Code82-002-4) and 1x EDTA-free Protease Inhibitor Cocktail (Thermo scientific, 1861279). Homogenates were passed through two layers of cheesecloth and centrifuged at 8,500g for 10 min to remove lipids in the supernatant and to pellet debris, nuclei and mitochondria. The pellet was resuspended in ice-cold HES buffer, transferred to a clean tube, and centrifuged at 700g for 10 min to spin down nuclei and cell debris. The supernatant was transferred to a clean tube again and centrifuged at 8,500g for 10 min to pellet mitochondria. The mitochondrial pellet was then washed with HES buffer twice and either pelleted for protein extraction or resuspended in RES buffer for mitochondrial HK1 activity assay or respiration assay.

Similar approach was used for mitochondria isolation from HEK 293 cells. Cultured cells were collected from culture dishes with approximately 80%-90% density of cells. The cell pellet was then homogenized with a motorized Teflon pestle. Homogenates were centrifuged at 700g for 5 min to pellet debris and nuclei. The supernatant was transferred to a clean tube again and centrifuged at 10,000g for 10 min to pellet mitochondria. The mitochondrial pellet was then washed with HES buffer twice and pelleted for protein extraction.

### Mitochondrial HK1 activity assay

Mitochondria HK1 activity was measured using the Glucose-6-Phosphate (Glc-6-P) dehydrogenase-coupled assay, coupling Glc-6-P formation to production of NADPH, monitored at absorbance of 340 nm by a plate reader (ThermoScientific, VARIOSKAN LUX), in the presence of excess Glc-6-P dehydrogenase (Sigma, G8519) and NADP+ (Roche, 10128031001). Isolated synaptosomal and BAT mitochondria suspended in RES buffer were used for mitochondrial HK1 activity assay.

For each synaptosomal mitochondrial HK1 activity reaction, 7.5 µg mitochondria suspension supplemented with 10 mM sodium pyruvate, 5 mM malate, 10 mM glucose, 100 µM ADP or supplemented with 10 mM glucose and 100 µM or 1 mM ATP was plated in a well of 96 well plate for continuous 30 min measurement (10s time intervals) at 37°C with constant medium shaking. For BAT mitochondrial HK1 activity reaction, 15 µg mitochondria suspension supplemented with 100 μM palmitate, 10 mM α-Ketoglutarate or 10 mM succinate, 10 mM glucose, 100 µM ADP was plated in a well of 96 well plate for continuous 30 min measurement (10s time intervals) at 37°C with constant medium shaking.

### Synaptosomal mitochondrial respiration assay and adenine nucleotide quantitation

Synaptosomal and BAT mitochondria respiration was measured by high-resolution respirometry with Oroboros Oxygraph-2k (Oroboros Instruments, Innsbruck, Austria) in a standard configuration, with 2 ml volume of the two chambers, at 37°C, and 700 rpm stirrer speed. For each measurement, 100 µg synaptosomal mitochondria or 200 µg BAT mitochondria suspended in 2 mL of RES were added to the respiration chamber. For synatosomal mitochondira, respiration was stimulated by injecting 10 mM pyruvate, 5 mM malate and 100 µM ADP with or without 10 mM glucose. For BAT mitochondria, respiration was stimulated by injecting 10 mM α-Ketoglutarate or 10 mM succinate. The software DatLab7 was used for data acquisition (2s time intervals) and analysis. All measurements were performed within 3 hours of mitochondria isolation.

For nucleotide quantitation, 100 µg synaptosomal mitochondria or 200 µg BAT mitochondria suspended in 2 mL of RES were added to the respiration chamber of Oroboros Oxygraph-2k with 700 rpm stirrer speed. For synaptosomal mitochondira, 10 mM pyruvate, 5 mM malate, and 100 µM ADP were injected. For BAT mitochondria, 10 mM α-Ketoglutarate or 10 mM succinate, and 100 µM ADP were injected. Samples were collected at indicated time points. ATP and ADP were extracted, processed and measured by the same approach as used with primary brown adipocytes.

### Proteomics analysis

Protein pellets were dissolved in 50 mM triethylammonium bicarbonate (TEAB, pH=8), 5% SDS at 60 °C with shaking for 30 min. Protein concentration was determined with a BCA method. Samples were then reduced by adding dithiothreitol (DTT) to a final concentration of 10 mM and samples were incubated at 56°C for 30 min. After cooling, iodoacetamide was added to a final concentration of 20 mM and samples were alkylated for 30 min at room temperature in the dark. Following centrifugation for 2 min at 13.2 krpm, the supernatants were removed and digested overnight with trypsin at 37°C using an S-Trap (Protifi). Following digestion, the peptide eluate was dried and reconstituted in 100 mM TEAB buffer. The samples were labelled with TMT10plex reagents, quenched with 5% hydroxylamine, and combined. For mitochondrial samples, the WT replicates were labeled with TMT labels 126, 127C, 127N, and 128 N, while the KO replicates were labeled with 128C, 129N, 129C, and 130N. The mixed samples were dried in a SpeedVac and reconstituted in a 2% acetonitrile, 0.1% TFA buffer to a final concentration of ∼0.5 ug/uL based on NanoDrop A205 (Thermo).

Samples were injected onto an Orbitrap Fusion Lumos mass spectrometer coupled to an Ultimate 3000 RSLC-Nano liquid chromatography system (Thermo). Samples were injected onto a 75 um i.d., 75-cm long EasySpray column (Thermo) and eluted with a gradient from 0-28% buffer B over 180 min. Buffer A contained 2% (v/v) ACN and 0.1% formic acid in water, and buffer B contained 80% (v/v) ACN, 10% (v/v) trifluoroethanol, and 0.1% formic acid in water. The mass spectrometer operated in positive ion mode with a source voltage of 1.8-2.0 kV and an ion transfer tube temperature of 300°C. MS scans were acquired at 120,000 resolution in the Orbitrap and top speed mode was used for SPS-MS3 analysis with a cycle time of 2.5 s. MS2 was performed with CID with a collision energy of 35%. The top 10 fragments were selected for MS3 fragmentation using HCD, with a collision energy of 55%. Dynamic exclusion was set for 25 s after an ion was selected for fragmentation.

Raw MS data files were analyzed using Proteome Discoverer v2.4 SP1 (Thermo), with peptide identification performed using a trypsin digest search with Sequest HT (cleavage after Lys and Arg except when followed by Pro). The mouse reviewed protein database from UniProt (downloaded Jan. 28, 2022, 17,062 entries) was used. Fragment and precursor tolerances of 10 ppm and 0.6 Da were specified, and three missed cleavages were allowed. A minimum peptide length of 6 residues was required. Carbamidomethylation of Cys and TMT10plex labelling of N-termini and Lys sidechains were set as a fixed modification, with oxidation of Met set as a variable modification. The false-discovery rate (FDR) cutoff was 1% for all peptides. At least two unique peptides were required for protein identification.

### Synaptosomal mitochondrial gold labeling

Synaptosomal mitochondrial gold labeling experiment was based on an earlier study (Kottke et al., PMID: 2457393). Mitochondria were suspended in isolation buffer without EGTA. 100 µL of the suspension containing 2 mg/ml of mitochondria protein were incubated for 10 min at room temperature with CEND1 (Cell signaling, 8944) or HK1 antibody (Invitrogen, MA5-14789) at a dilution of 1:100. The mitochondria were washed three times and resuspended in 100 µL isolation buffer and incubated with 3 µL protein-A/Gold (Electron Microscopy Sciences, 25284) for 10 min at room temperature. The samples were centrifuged, washed and resuspended in a buffer containing 125 mM KCl and 10mM HEPES for cryo-ET or subjected to fixation with 2.5% glutaraldehyde in 0.1 M cacodylate buffer (pH 7.4) for EM analysis.

### Cryo-ET analysis

The freshly isolated mouse brain synaptosomal mitochondria labeled with CEND1 antibody and gold-labeled protein A were diluted with PBS to a final protein concentration of 0.1mg/ml. Next, 3.5µl of the diluted mitochondria sample was applied to a Lacey carbon 300 mesh copper grid (Ted Pella). The grid was pre-treated using a PELCo easiGlow instrument (Ted Pella) for 80s with 30mA current. The grid was backblotted using a Leica EM GP2 (Leica) for 7s with the sensor turned on before plunging into liquid ethane for vitrification.

Grid imaging was performed on a Titan Krios G3 microscope (Thermo Fisher Scientific) equipped with a Falcon 4i camera (Thermo Fisher Scientific). Tilt series were collected using the PACEtomo (PMID: 36456783) script in SerialEM (PMID: 16182563). Dose-symmetric scheme with increments of 3° in groups of two tilts (PMID: 27313000) was applied. Tilt range was set between -60° and +60°, resulting in a total number of 41 tilt images per tilt series. The total dose per tilt image was approximately 3e-/Å2.

Tilt series data processing was performed using the tomography pipeline in Relion 5 (PMID: 39147729). Briefly, motion correction and dose weighting was done using the implementation within Relion (PMID: 30412051); odd/even tilt series were calculated for subsequent denoising jobs. Next, dark/empty images were manually removed using the “Exclude tilt-images” in Relion 5 (PMID: 39147729). Tilt series alignment was performed using the AreTomo2 (PMID: 35601683; https://github.com/czimaginginstitute/AreTomo2) wrapper in Relion 5. CTF (contrast transfer function) of the images was also estimated using AreTomo2 during the tilt seriels alignment process. Tomograms were then reconstructed using Relion 5, generating odd/even tomograms for denoising. Tomogram denoising was performed using the Relion 5 wrapper to cryo-CARE (Buchholz T-O, Jordan M, Pigino G and Jug F(2018), Cryo-CARE: content-aware image restoration for Cryo-transmission electron microscopy data. DOI: 10.1109/ISBI.2019.8759519) with default settings. Tomogram visualization was done using UCSF ChimeraX (PMID: 32881101).

#### Acknowledgements

We thank Andrew Lemoff and the Proteomics core at UTSW for the proteomics analysis; the EM core at UTSW (supported by NIH grant 1S10OD021685-01A1), especially Dr. Natalia Gunko, Phoebe Doss, and Ethan Ozment, for project discussion and EM sample preparation. X.Z. is a Rita C. and William P. Clements, Jr. Scholar in Biomedical Research and was supported by the Endowed Scholars in Medical Science Program at UTSW, Cancer Prevention and Research Institute of Texas grant RR200084, NIH grant R01DK135556, Welch Foundation Award I-2220-20240404, and American Heart Association Career Development Award 23CDA1050474. J.R.G. and K.M.H. were supported by NIH grant U54HD104461 and K.M.H. was supported by R37NS114516. B.W. is a Southwestern Medical Foundation Scholar in Biomedical Research and supported by NIH grant R35GM160160 and a Research Grant from Robert Welch Foundation (I-2171). R.M. was supported by Hellenic Foundation for Research and Innovation Project Number HFRI-FM17C3-1019. Advance Imaging Research Center at UT Southwestern was partially funded by the Cancer Prevention and Research Institute of Texas grant RP210099. Cryo-ET studies at the Structural Biology Laboratory and the Cryo-Electron Microscopy Facility at UT Southwestern Medical Center were partially supported by the Cancer Prevention & Research Institute of Texas grant RP220582.

## Author contributions

X.Z. conceived the project. X.Z. and D.Y. designed the experiments. D.Y., C.Z., D.P., M.B., and M.L. performed the majority of the experimental work. G.M., N.N., J.G., and K.M. conducted the electrophysiological studies. Y.H. carried out cryo-ET data collection and processing. X.W., S.P., and J.W. performed the PET-MRI imaging. B.W. contributed to nucleotide-analysis method development. H.B. conducted LC/MS-based nucleotide analysis. M.G. and R.M. provided the *Cend1* KO mice. X.Z. and D.Y. wrote the manuscript, and all authors contributed to manuscript review and discussion.

## Competing interests

The authors declare no competing interests.

## Materials & Correspondence

Correspondence and material requests should be addressed to Xing Zeng at.

---

**Figure S1.**
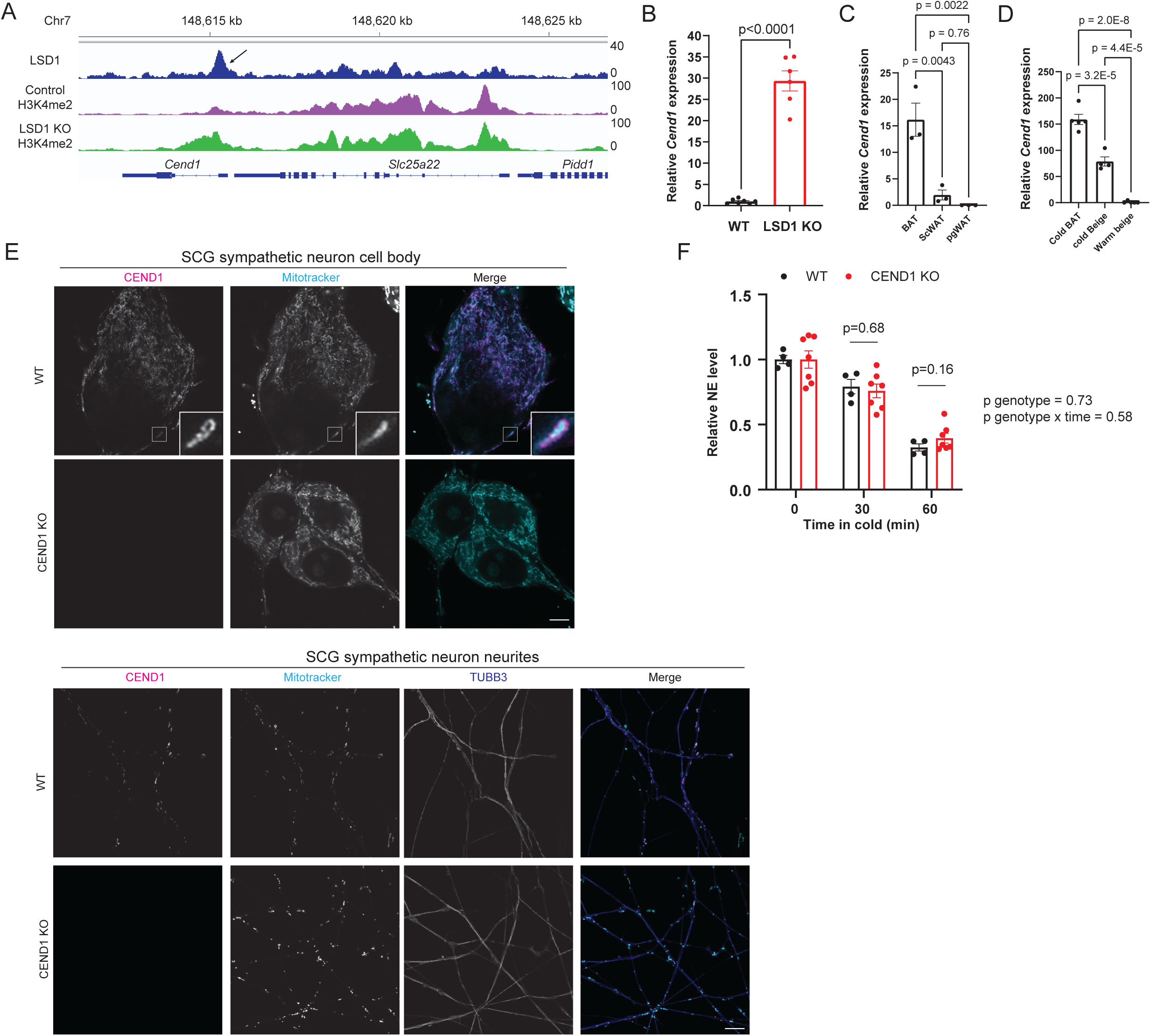
A, brown adipose tissue LSD1 and H3K4me2 ChIPseq reads distribution at the *Cend1* locus. Note the markedly increased H3K4me2 signal near the LSD1 binding site at the *Cend1* locus in the LSD1-deficient brown adipose tissue. B, qPCR analysis of *Cend1* expression in control or adipose-specific LSD1 KO brown adipose tissue. C, *adq-cre* driven TRAP-seq analysis of Cend1 expression in adipocytes of various depots at RT. D, *ucp1-cre* driven TRAP-seq analysis of Cend1 expression in brown or beige adipocytes. E, immunofluorescence analysis of CEND1 expression in WT and CEND1 KO primary superior cervical ganglia (SCG) sympathetic neurons. The inset shows CEND1 staining pattern consistent with OMM localization. Scale bar: 5 µm. F, NE turnover rate in WT and CEND1 KO brown adipose tissue after acute cold exposure and a-methyl-*p*-tyrosine-methyl ester injection. Statistical analysis was performed with unpaired t-test in B, one-way ANOVA with tukey in C and D, and two-way ANOVA with repeated measurement in F.

**Figure S2.**
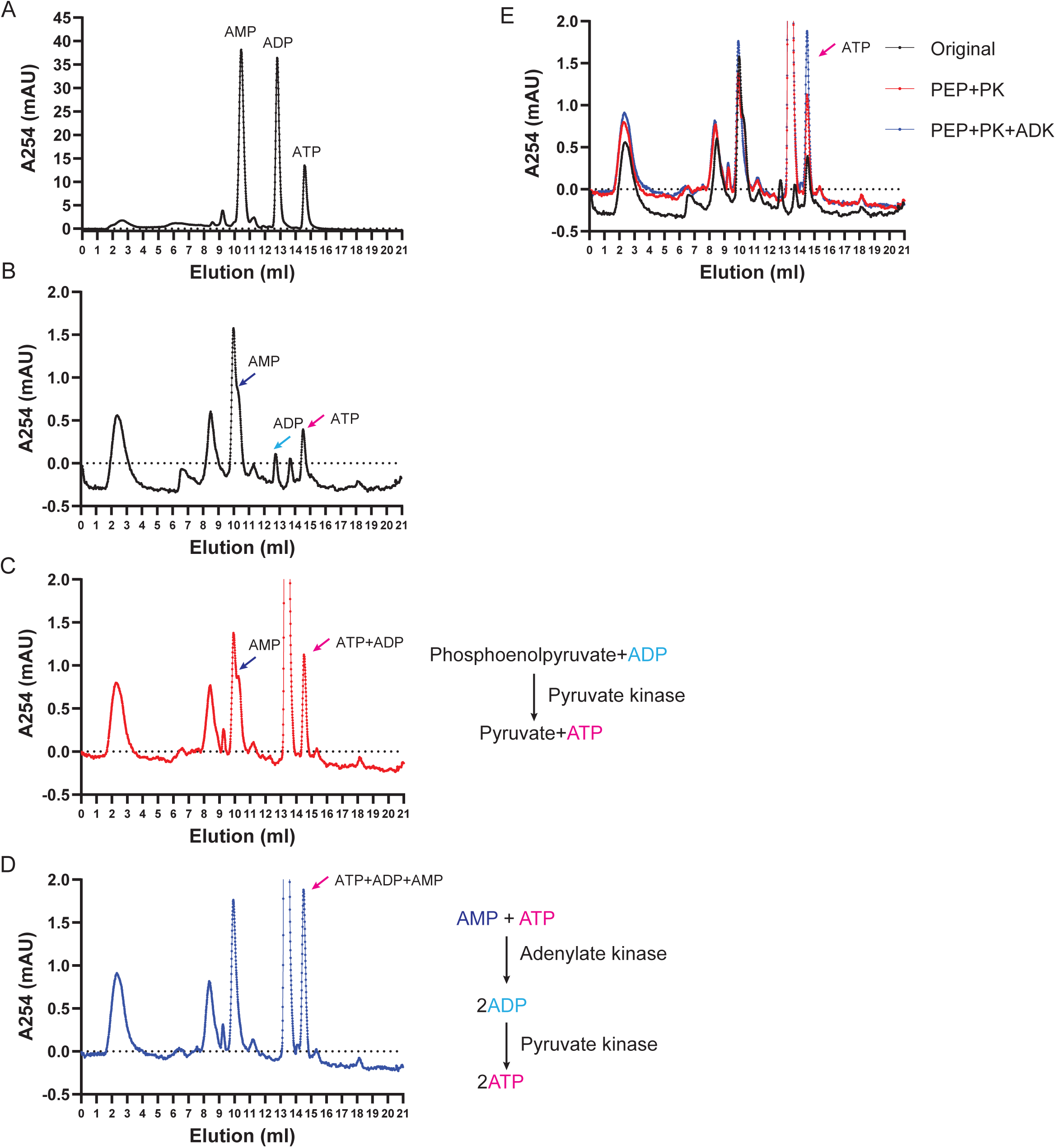
A, elution profile of ATP, ADP, and AMP standards from CaptoHiRes Q column monitored by absorbance at 254 nm. B, extract from brown adipocytes stimulated with NE. Note the ATP, ADP, and AMP peaks. The AMP peak is on the shoulder of an uncharacterized metabolite. C, conversion of ADP into ATP by the addition of PEP and pyruvate. Note the disappearance of the ADP peak and the increase in the ATP peak. D, further conversion of AMP into ATP by the addition of adenylate kinase. Note the disappearance of the AMP peak and the increase in the ATP peak. E, overlay of the traces to show the stepwise increase in the ATP peak, validating this method for ATP/ADP/AMP quantitation.

**Figure S3.**
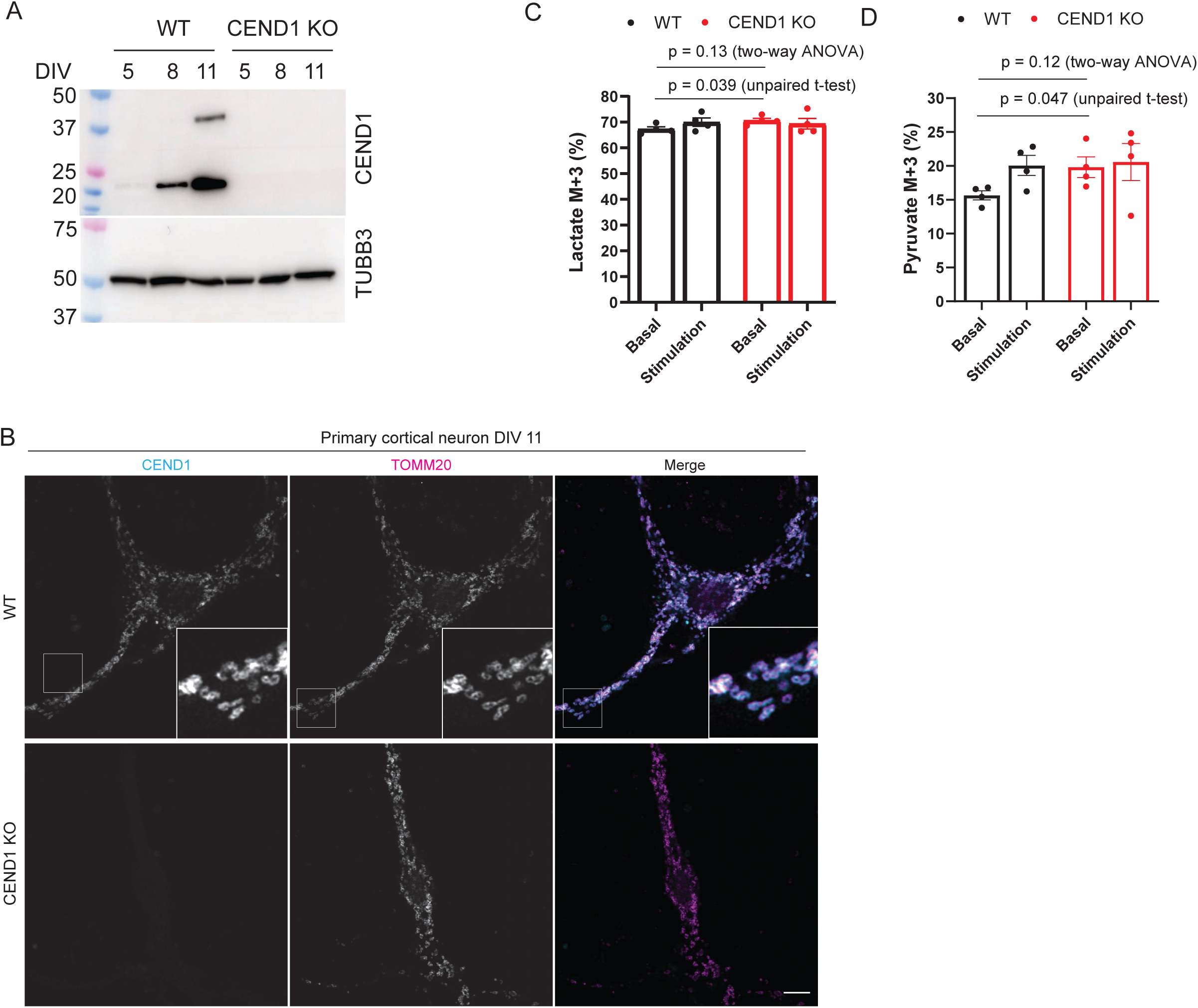
A, western blot analysis of CEND1 expression in primary cortical neurons cultured *in vitro*. Note the time-dependent increase in CEND1 expression as neurons mature. B, immunofluorescence microscopy analysis of CEND1 expression in primary cortical neurons cultured *in vitro* on DIV11. Scale bar: 5 µm. C-D,^13^C labeling efficiency of lactate (C) or pyruvate (D) in DIV11 primary WT and CEND1 KO cortical neurons with or without electrical stimulation. Statistical significance between WT and CEND1 KO neurons under the resting condition was tested with unpaired t-test or two-way ANOVA as indicated in the figure.

**Figure S4.**
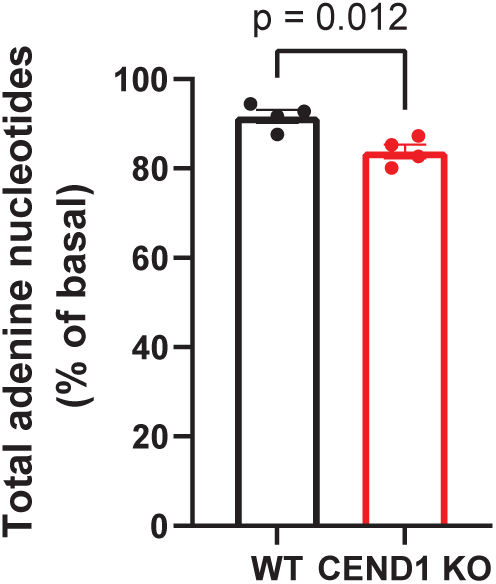
NE stimulation-induced relative reduction in total adenine nucleotides in primary WT and CEND1 KO brown adipocytes. Statistical significance was calculated by unpaired t-test.

**Figure S5.**
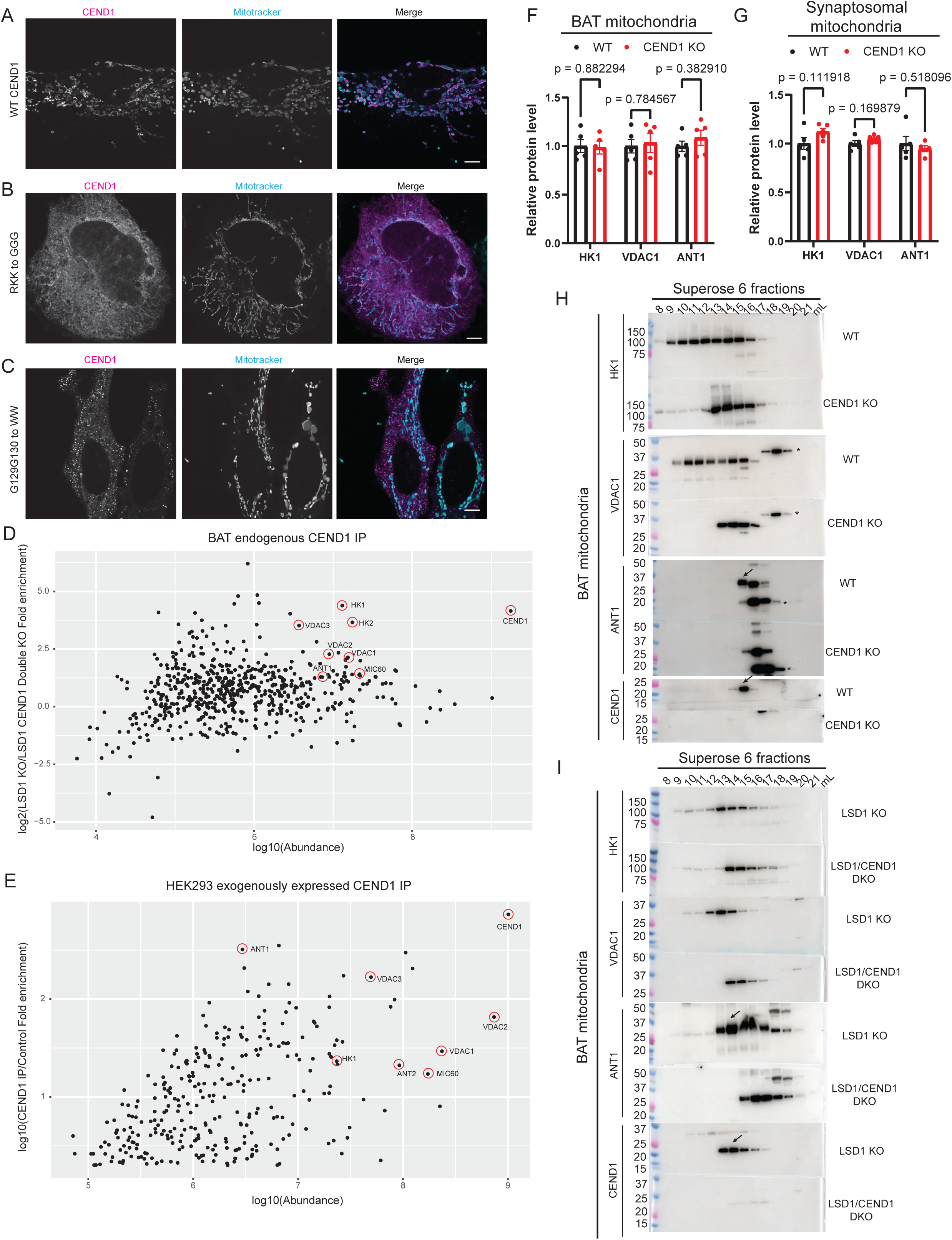
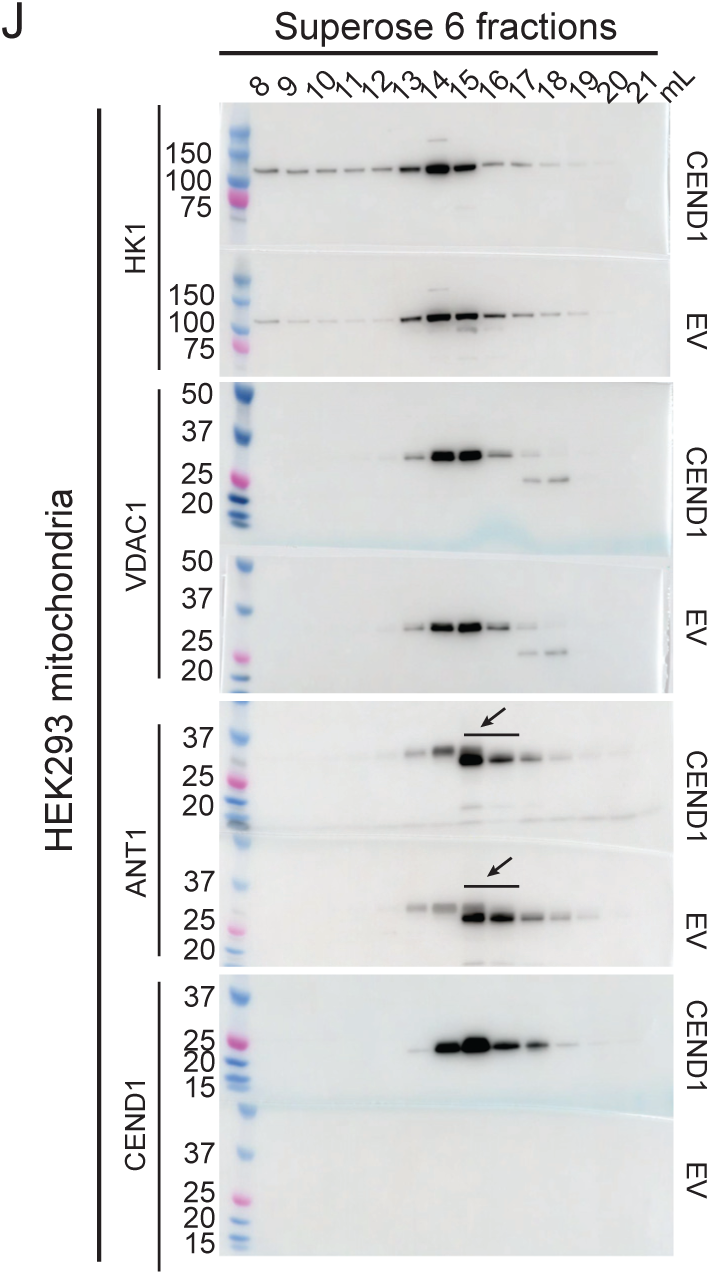
A-C, immunofluorescence microscopy of subcellular localization of WT CEND1 (A), RKK to GGG (B), or G129G130/WW mutants. D, mass spectrometry analysis of proteins isolated with the CEND1 antibody from LSD1 KO BAT mitochondria or LSD1/CEND1 double KO BAT mitochondria. For each protein, the LSD1 KO/LSD1 CEND1 double KO fold enrichment is plotted against the abundance. E, mass spectrometry analysis of proteins isolated with the CEND1 antibody from HEK293 cells transfected with a CEND1-expressing vector or an empty vector. For each protein, the LSD1 KO/LSD1 CEND1 double KO fold enrichment is plotted against the abundance. A 2-fold enrichment cutoff is applied to avoid cluttering. F, protein abundance quantitated by proteomic analysis of WT or CEND1 KO BAT mitochondria. G, protein abundance quantitated by proteomic analysis of WT or CEND1 KO synaptosomal mitochondria. H-J, gel filtration analysis of mitochondrial extract HK1/VDAC1/ANT1/CEND1 elution profiles prepared from WT or CEND1 KO mouse BAT (H), LSD1 KO or LSD1/CEND1 double KO mouse BAT (I), or HEK293 cells transfected with a CEND1-expressing or empty vector (J).

**Figure S6.**
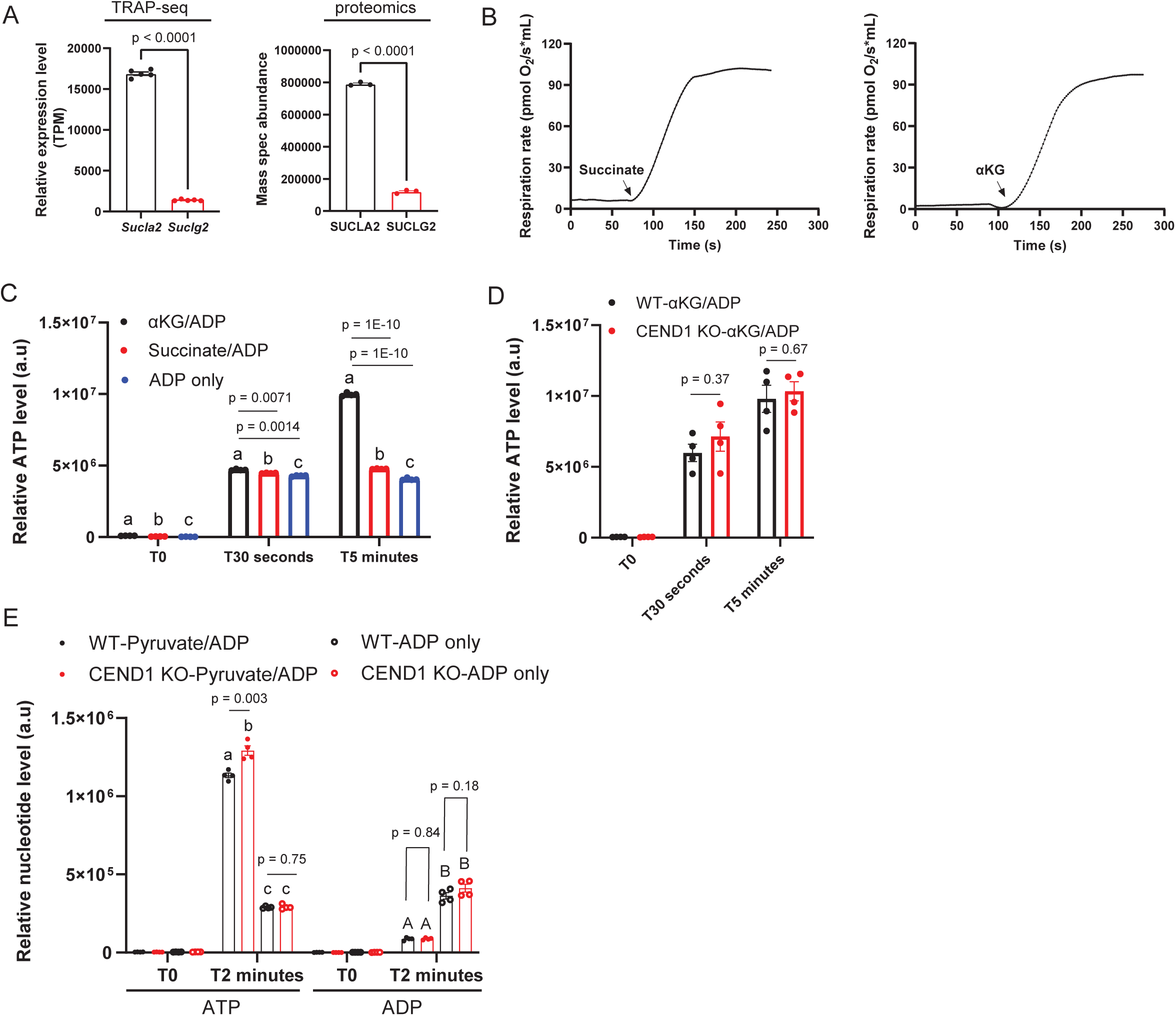
A, relative expression level of *Sucla2* and *Suclg2* in brown adipocytes detected by TRAP-seq or proteomics. B, representative oxygen consumption rate traces from brown adipocyte mitochondria supplemented with 5 mM succinate or a-ketoglutarate. C, ATP quantitation from brown adipocyte mitochondria respiring on 5 mM succinate or a-ketoglutarate or only supplemented with 100 µM ADP. Note the conversion of a fraction of ADP into ATP by adenylate kinase in the absence of any oxidizable substrate. E, ATP/ADP quantitation from synaptosomal mitochondria supplemented with 100 µM ADP with or without 5 mM pyruvate/malate. Note the conversion of a fraction of ADP into ATP by adenylate kinase in the absence of any pyruvate/malate. Statistical significance was calculated by unpaired t-test in A, D and E, and one-way ANOVA with Tukey in C.

## Notes

### Competing Interest Statement

The authors have declared no competing interest.

